# Robust regulatory interplay of enhancers, facilitators, and promoters in a native chromatin context

**DOI:** 10.1101/2025.07.07.663560

**Authors:** Zhou Zhou, Albert Li, Junke Zhang, Haiyuan Yu, Abdullah Ozer, John T. Lis

## Abstract

Enhancers are gene-distal *cis*-regulatory elements that drive cell type-specific gene expression. While significant progress has been made in identifying enhancers and characterizing their epigenomic features, much less effort has been devoted to elucidating mechanistic interactions among clusters of functionally linked regulatory elements within their endogenous chromatin contexts. Here, we developed a novel recombinase-mediated genome rewriting platform and applied our divergent transcription architectural model to understand how a long-range human enhancer confers a remarkable 10,000-fold activation to its target gene, *NMU,* at its native locus. Our systematic dissection reveals transcription factor synergy at this enhancer and highlights the interplay between a divergently transcribed core enhancer unit and emerging new types of *cis*-regulatory elements—notably, intrinsically inactive facilitators that augment and buffer core enhancer activity, and an adjacent retroviral long terminal repeat promoter that represses enhancer activity. We discuss the broader implications of our focused study on enhancer mechanisms and regulation genome-wide.

## Introduction

Since their discovery over four decades ago,^1,2^ enhancers have been recognized as abundant and essential *cis*-regulatory elements that recruit transcription factors (TFs) to activate target gene promoters from a distance, often in a cell type-specific manner. Owing to their pivotal roles in development and disease, numerous individual laboratories and major consortia like ENCODE^3^ have made extensive efforts to identify and characterize enhancers across diverse cell types and tissues. Traditional hallmarks of active enhancers include TF and coactivator binding, DNase I Hypersensitivity, and histone modifications such as H3K27ac, H3K4me1 and H3K4me3.^4^ Later, widespread RNA Polymerase II (Pol II) transcription has emerged as another key indicator of enhancer activity.^5^ We previously developed the nuclear run-on–based assays, GRO-cap and PRO-cap,^6,7^ which selectively enrich for 5′-capped nascent RNAs to map genome-wide transcription initiation events with high sensitivity and specificity. Applying GRO-cap to human cells revealed a unified molecular architecture shared by enhancers and promoters, featuring a central nucleosome-depleted TF binding region flanked by two divergent core promoters that initiate bidirectional Pol II transcription.^8^ This unit definition enables precise delineation of enhancer boundaries and offers a robust framework for accurate enhancer annotation.^9,10^ However, functional dissection of enhancers within the paradigm of divergent transcription architecture remains limited.

Beyond the correlative features, recent technological advancements have further established two powerful types of high-throughput screening methods to directly quantify enhancer activity. Gain-of-function assays, such as massively parallel reporter assays (MPRAs)^11,12^ and self-transcribing active regulatory region sequencing (STARR-seq),^13^ measure elements’ intrinsic enhancer potential based on their ability to drive reporter gene transcription. While these assays allow impressive genome-wide scalability, they rely on assaying DNA sequences outside of their endogenous chromatin contexts, which may compromise physiological relevance and introduce false positives.^13^ In contrast, CRISPR-based loss-of-function screens^14–16^ assess enhancer necessity and preserve the native spatial relationships between enhancers and promoters, but they are complicated by variable perturbation efficiency, potential off-target effects, and imprecise definition of element boundaries. Moreover, human genes are frequently regulated by ensembles of enhancers and related elements that can act redundantly,^17^ additively, or synergistically,^18–21^ representing additional layers of complexity. Therefore, despite the exponential rise in the number of experimentally nominated *cis*-regulatory elements, the mechanisms governing their functional logic are still poorly understood.

To address these limitations and provide orthogonal insights, recombinase-mediated genome rewriting^22–24^ has emerged as a powerful strategy. Through precise replacement of genomic regions and targeted manipulation of individual or combinatorial elements, these approaches allow comprehensive interrogation of entire loci of interest and enable functional analysis in their native genomic contexts, uncovering hierarchical relationships among a cluster of elements. Furthermore, they embrace serendipitous discovery of previously unrecognized *cis*-regulatory behavior. For instance, by engineering the *ɑ-globin* super-enhancer in mouse erythroid cells, Blayney et al.^24^ recently identified a novel class of distal regulatory elements, termed **facilitators**, which lack intrinsic enhancer activity but potentiate the function of autonomous enhancers. While conceptually intriguing, the broader prevalence of facilitators beyond super-enhancers and the molecular underpinnings of enhancer– facilitator interactions remain largely unexplored.

In this study, we developed a novel recombinase-mediated platform to systematically dissect a potent distal human enhancer at its native locus, guided by our architectural model of enhancer organization.^8,9^ Through detailed TF motif mutagenesis and integrative genomics analysis, we uncovered intricate crosstalk at both the *trans*-acting factor and the *cis*-acting element levels. We demonstrate that a core enhancer region, precisely demarcated by a divergent transcription pattern, acts as the intrinsic activating unit for target gene expression. This core enhancer activity is further modulated by surrounding facilitators and a promoter-like element, which display distinct molecular signatures and exert positive and negative influences, respectively. We propose that such highly interconnected regulatory networks are broadly utilized across the genome to ensure precise and robust control of transcriptional output.

## Results

### eNMU landing pad as a powerful system to study enhancer function at the native locus

Neuromedin U (NMU) is a neuropeptide that has been implicated in various physiological processes including erythropoiesis.^25^ In the triploid human erythroleukemia cell line K562, Gasperini et al.^15^ identified a critical enhancer of *NMU* (hereafter eNMU) in a CRISPR interference (CRISPRi) screen. This enhancer is located ∼94 kb away from the *NMU* gene promoter, and its homozygous deletion, without negatively affecting cell growth, led to a remarkable 100% reduction in *NMU* expression by RNA-seq.^15^ Tippens and Liang et al.^9^ further refined the boundary of eNMU based on our unified molecular architecture model of transcriptional regulatory elements^8,26^ and divided eNMU into two divergently transcribed sub-elements e1 and e2 (Figure 1A). Homozygous CRISPR knockouts showed that deleting e1, the 453-bp sub-element with higher DNase I Hypersensitivity (DHS), reduced gene expression by 10,000-fold (0.01% of WT) by quantitative reverse transcription PCR (RT-qPCR)—the same level as deleting full eNMU (Figure 1B). Precision Run-On and Sequencing (PRO-seq) confirmed that ΔeNMU and Δe1 abolished nascent transcription at both the enhancer and the target gene without affecting other genes nearby (Figures 1C, 1D, and S1). Hence, e1 is essential for transcription initiation while e2 alone in the genome is completely inactive. Surprisingly, deletion of this intrinsically inactive 503-bp e2 element resulted in only ∼5% of WT mRNA level (Figure 1B) with decreased PRO-seq signal at e1 and the *NMU* gene, highlighting that e1 acts as a canonical autonomous enhancer for *NMU* but requires the **facilitator** element^24^ e2 to achieve maximal activation.

**Figure 1.**
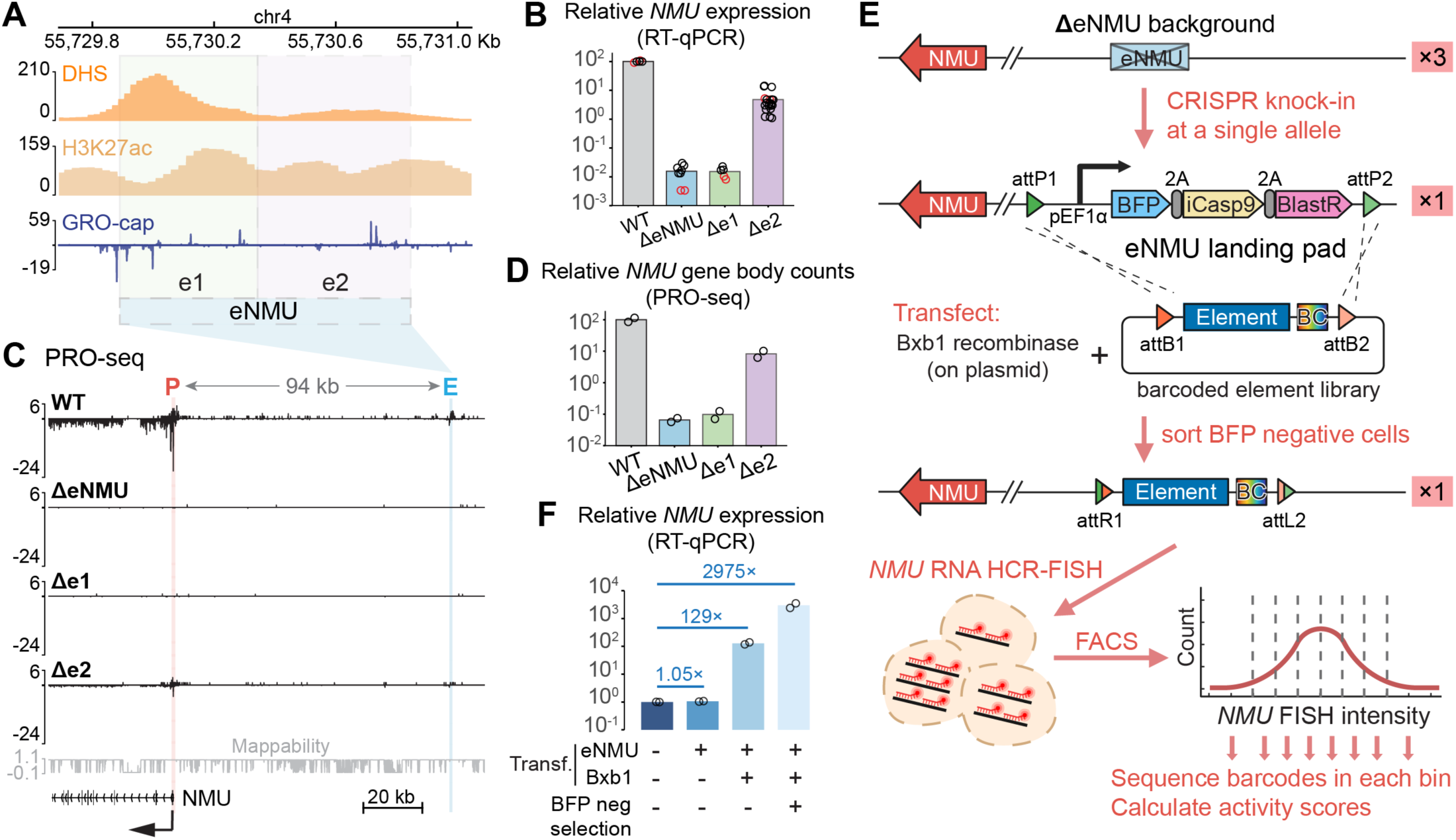
eNMU landing pad as a powerful system to study enhancer function at the native locus. (A) Epigenomic landscape of eNMU and its sub-elements e1 and e2 in K562 cells, showing DNase I hypersensitivity (DHS),^3^ H3K27ac,^3^ and GRO-cap–defined transcription start sites (TSSs).^8^ (B) *NMU* mRNA levels (RT-qPCR) in independent cultures of WT, ΔeNMU, Δe1 and Δe2 cell lines. Black dots = data from Tippens and Liang et al.^9^ (*GAPDH* normalized); red dots = data from this study (*ACTB* normalized). (C) PRO-seq signal at the *NMU*–eNMU locus in the same cell lines as (B). Tracks represent merged biological replicates (n = 2). (D) Relative *NMU* gene body read counts from PRO-seq in (C). (E) Workflow of the eNMU landing pad system to measure enhancer activity of a barcoded element library. (F) Rescue of *NMU* expression by inserting eNMU into the landing pad (LP); n = 2 independent LP clones. See also Figures S1 and S2.

The huge dynamic range of eNMU regulation from a distal site and the intriguing cooperativity between its sub-elements e1 and e2 warrant a comprehensive interrogation into its sequence features and molecular mechanisms. To this end, we used CRISPR to knock in a landing pad at a single allele of the native eNMU locus in the K562 ΔeNMU cell line (Figure 1E and S2A). This landing pad, modified from Matreyek et al.,^27^ harbors a Bxb1 recombinase-mediated exchange cassette containing a battery of selection markers, including a constitutively expressed Blue Fluorescent Protein (BFP). Co-transfection of a Bxb1-expressing plasmid and a barcoded payload plasmid library of eNMU mutants leads to the loss of selection markers and a stable, irreversible integration of individual elements at the landing pad locus. The resulting BFP^−^ recombinant population is then subjected to *NMU* hybridization chain reaction fluorescence *in situ* hybridization coupled with flow cytometry (HCR-FlowFISH)^28^ to resolve the effects of individual mutants on *NMU* expression as a measure of enhancer activity. As an initial test on the functionality of our system, we integrated the full-length eNMU sequence into two independently isolated landing pad (LP) clones and observed ∼5% recombination efficiency in both cases as measured by BFP loss (Figure S2B). Importantly, for both LP clones, *NMU* expression was rescued by ∼3,000-fold in BFP^−^ recombinant cells compared to the parental, *NMU*-inactive LP cells (Figure 1F and S1), consistent with 1/3 of the 10,000-fold activation by three alleles of eNMU (Figure 1B). Subsequent testing of HCR-FlowFISH on eNMU- and e2-recombinant populations showed a clear separation of their *NMU* RNA FISH signals (Figure S2C). Therefore, we have successfully established an efficient LP-based workflow that would allow us to characterize eNMU in its native chromatin context.

### Screening for functional units and motifs in eNMU

We next set out to design a systematic mutagenesis scheme for eNMU using a combination of unbiased and targeted perturbations that complement each other. Building on our previous finding that enhancer transcription contributes to activity,^9^ we generated tiling deletions for e1 and e2 to remove clusters of transcription start sites (TSSs) (Figure 2A). In parallel, considering the central role of TFs in shaping enhancer function, we curated a list of TF motifs for targeted mutagenesis by (1) intersecting all possible TF binding sites (TFBSs) from the JASPAR database^29^ with K562 ChIP-seq peaks from the UCSC Genome Browser^30^ overlapping eNMU, and (2) filtering this candidate set to retain only those motifs corresponding to K562-expressed TFs and located within regions of enriched ChIP-seq signals (Methods). This led to a total of 95 motifs corresponding to 26 TFs (Figure S3A). We then merged highly similar motifs such as AP-1 and NFE2 and mutated the two most conserved bases in each motif by transversion^31–33^ (A↔C, T↔G, Table S1) while ensuring minimal interference with overlapping motifs to the best of our ability. Finally, we devised a “mix-and-match” cloning strategy (Figure S2D; Methods) to construct a barcoded mutant library where all the motif occurrences for a given TF were altered only in e1, e2, or both. This approach allowed for maximal disruption of TF binding within the distinct contexts of the two sub-elements. Taken together, our library contains 83 elements (77 eNMU-derived sequences and 6 exogenous controls) associated with 328 unique barcodes—on average 4 barcodes per element (Tables S1 and S3). Following library integration and recombinant cell selection, we performed FlowFISH and sorted cells into 8 bins based on *NMU* signal intensity, using *ACTB* as an internal control (Figure S2E). We then sequenced enhancer barcodes in each bin and calculated an activity score for each barcode using a weighted average approach (Figures S2F and S2G). Activity scores from biological replicates showed a strong correlation (Pearson’s r = 0.91, Figure S2H) and aligned closely with RT-qPCR measurements for a select set of mutants (Pearson’s r = 0.97, Figure S2I), confirming that our assay faithfully captured mutant activities.

**Figure 2.**
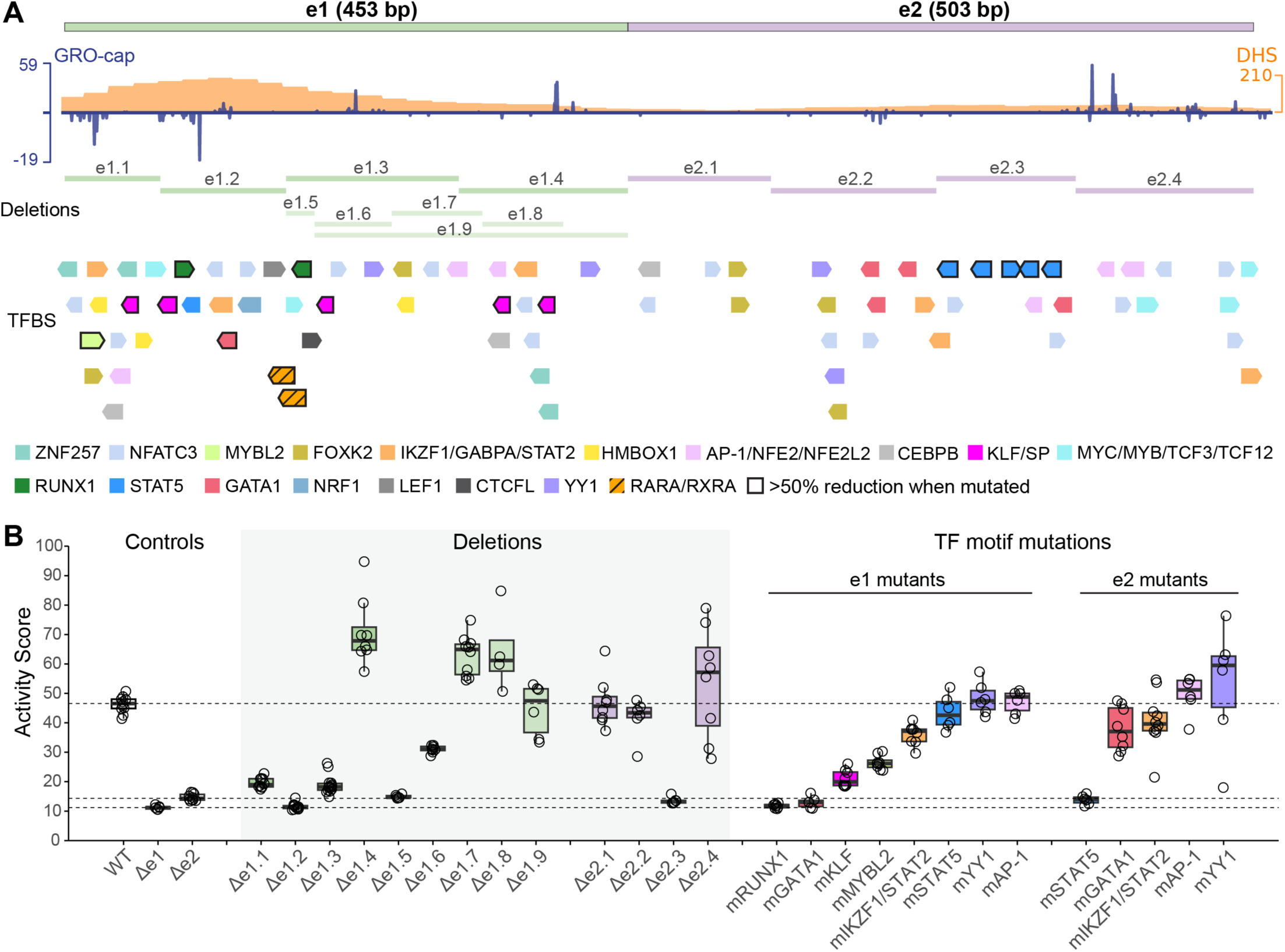
Screening for functional units and motifs in eNMU. (A) Overview of deletions and targeted TF binding sites (TFBS) in the eNMU mutagenesis screen. (B) Activity scores of select mutants measured by HCR-FlowFISH. Each dot represents the score of an element-specific barcode from either of the two biological replicates. Dashed lines indicate median scores of control elements WT_eNMU, Δe1, and Δe2. See also Figures S2 and S3; Tables S1 and S3.

Our analysis of tiling deletions in e1 showed that a divergently transcribed region e1.1–e1.3 delineated an activating unit, where e1.2—encompassing the DHS signal summit—marked the core of e1 activity (Figure 2B, Δe1.2 versus Δe1). Unexpectedly, the well transcribed e1.4 acted as a repressing unit, as its deletion led to an increase in *NMU* expression. This observation was bolstered by finer deletions Δe1.5 to Δe1.8, which revealed that the transition between activation and repression lay between e1.6 and e1.7. In fact, the first 201 bp of e1 (Δe1.9) was sufficient to capture all of its activity, likely due to the loss of both positive and negative elements in e1.9. In contrast, tiling deletions in the facilitator e2 identified a simpler functional core e2.3 (Figure 2B, Δe2.3 versus Δe2), with the other segments showing modest effects. Overall, fold changes in double deletions of an e1 segment and an e2 segment were multiplicative (i.e., log-additive) of single deletions, except for Δe1.2+Δe2.3, which fell below the dynamic range of FlowFISH (Figures S3B and S3D). These findings highlight a modular nature of eNMU’s molecular architecture with largely independent activating and repressing features.

Examination of the motif mutagenesis results in e1 or e2 showed <50% reduction of enhancer activity in most cases (Figures 2 and S3C). Consistent with the deletion results, the key TF motifs for e1, namely GATA1 and RUNX1 motifs, are located within or near the core region e1.2, while the essential motifs for e2—the STAT5 motifs—are clustered in the core e2.3, accounting for nearly all its function. TF contributions were context-specific, as exemplified by GATA1 being much more critical for e1 than for e2. Similarly, double mutants where the same TF binding was disrupted in both e1 and e2 exhibited multiplicative effects (Figures S3C and S3D), suggesting that the cooperativity between e1 and e2 is not driven by a single TF type but likely involves multiple different TFs.

### Interplay of regulatory factor binding at eNMU

To validate the findings on the key motifs and enable clean functional analysis downstream, we generated single cell recombinant clones harboring WT_eNMU, e1_mRUNX1, e1_mGATA1, and e2_mSTAT5. RT-qPCR quantifications of *NMU* expression showed marked reductions in the mutant clonal lines, corroborating our FlowFISH results (Figure 3A). Individual mutation of the left and right RUNX1 motifs revealed that the two sites acted additively, with a stronger contribution from the left site, consistent with its higher motif score (JASPAR scores 632 versus 335, Table S1). Substituting the GATA1 motif with a RUNX1 motif (e1_G-to-R) failed to rescue e1_mGATA1’s phenotype, suggesting GATA1 as an indispensable factor for eNMU function. Conversely, replacing the RUNX1 motifs with GATA1 (e1_R-to-G) only partially rescued e1_mRUNX1’s phenotype, suggesting that the combination of GATA1 and RUNX1 is particularly potent in the context of e1. We also noticed that the modest effect of mutating the right RUNX1 motif—located in the segment e1.5—was not sufficient to explain the substantial decrease in enhancer activity in the Δe1.5 mutant (Figure 2). A closer examination of this region uncovered two strong Retinoic Acid Receptor Alpha (RARA)/Retinoid X Receptor Alpha (RXRA) motifs (Figure 2A, orange hatched boxes), which overlapped the right RUNX1 motif and were initially overlooked due to the absence of publicly available ChIP-seq data. Mutating both RARA/RXRA motifs without disrupting the RUNX1 site validated their crucial function (Figure 3A, e1_mRARA). Therefore, we have established that four distinct TF motifs, including e1’s RUNX1, GATA1, and RARA/RXRA motifs and e2’s STAT5 motifs, are pivotal for eNMU activity.

**Figure 3.**
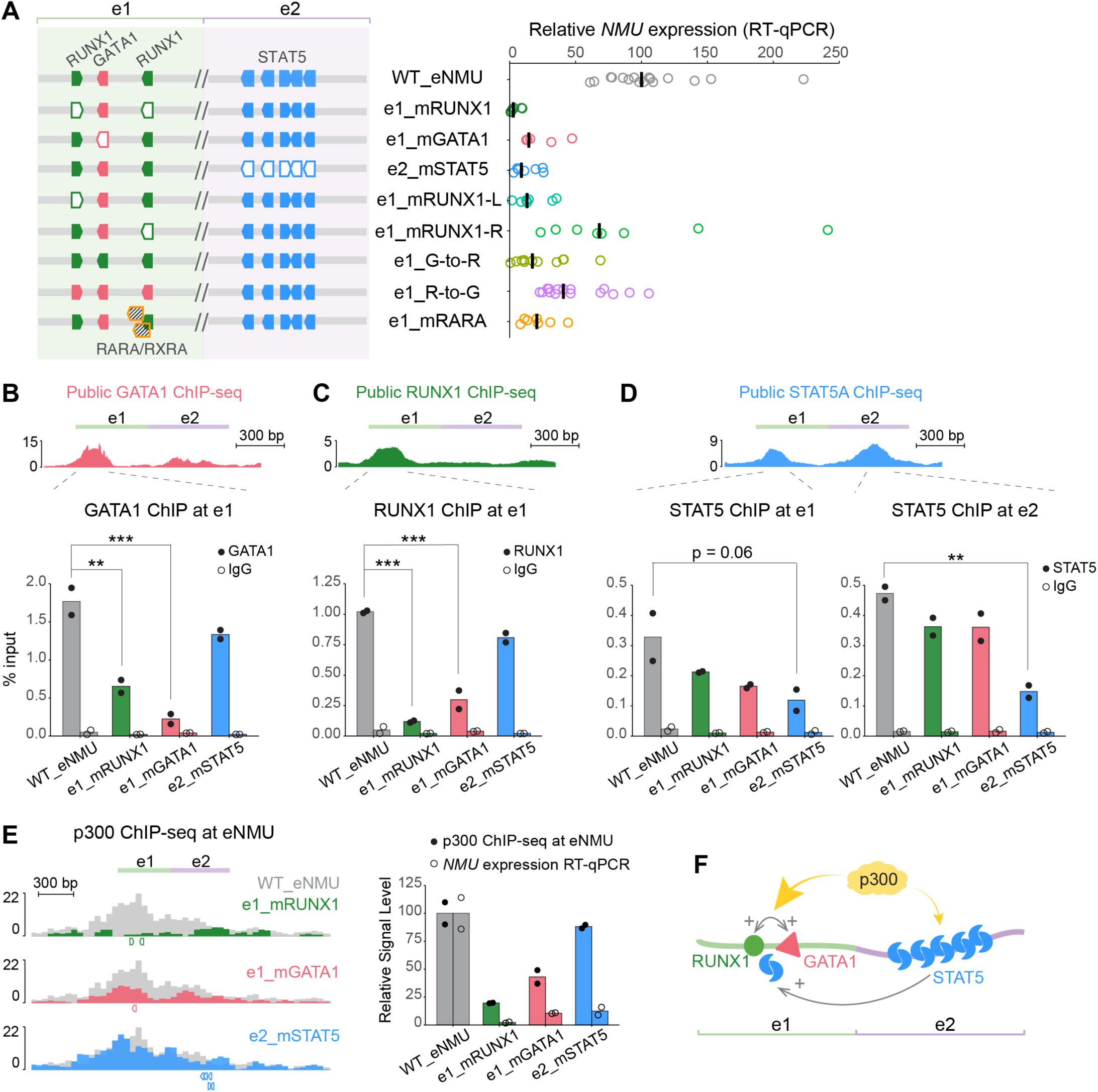
Interplay of regulatory factor binding at eNMU. (A) *NMU* mRNA levels measured by RT-qPCR in single cell-derived recombinant clones; bars = median. Exact mutant sequences are listed in Table S1. (B–D) ChIP-qPCR of TF binding in select mutants: GATA1 (B) and RUNX1 (C) at e1; STAT5 (D) at e1 and e2 (n = 2 independent single cell clones per mutant). Statistical significance assessed using one-way ANOVA with Dunnett’s post hoc test vs. WT_eNMU (**, p < 0.01; ***, p < 0.001). Public ENCODE ChIP-seq tracks (fold change over control)^3^ shown as references: GATA1, ENCFF334KVR; RUNX1, ENCFF654QOE; STAT5A, ENCFF171KLX. (E) Left: p300 ChIP-seq profiles at the eNMU locus in the indicated mutants from (B–D); tracks represent merged biological replicates (n = 2). Track colors indicate specific motif disruptions: grey = WT_eNMU, green = e1_mRUNX1, red = e1_mGATA1, blue = e2_mSTAT5. Colored boxes below tracks indicate locations of disrupted TF motifs. Right: p300 signal at eNMU vs. *NMU* mRNA in matched single cell clones. (F) Schematic of regulatory factor interplay at eNMU. Note that, unlike normal physiological conditions where STAT5 proteins are activated in response to cytokine signaling,^37^ K562 cells express the constitutively active oncogenic BCR-ABL fusion protein that drives persistent STAT5 phosphorylation, dimerization and activation.^38,39^ See also Methods.

We next investigated how the motif alterations functionally affected TF occupancy. ChIP-qPCR assays revealed that GATA1 and RUNX1 binding at e1 was significantly impaired not only by disruption of their own motifs, but also by mutations in each other’s motifs (Figures 3B and 3C), indicating cooperative binding of these two factors. STAT5 binding at e2 was largely self-driven, as demonstrated by its significant reduction in the e2_mSTAT5 mutant but minor changes in the e1 mutants (Figure 3D, right panel). Interestingly, STAT5 binding at e1 was also affected by e2’s STAT5 mutations (Figure 3D, left panel), suggesting that the facilitator element e2 might boost e1’s activity by promoting STAT5 binding at e1. Collectively, these results demonstrate extensive synergy in TF occupancy at eNMU (Figure 3F).

Since p300 is a known coactivator for GATA1,^34^ RUNX1,^35^ and STAT5,^36^ we also performed p300 ChIP-seq to examine its recruitment in the mutant clones. The RUNX1 and GATA1 mutants exhibited prominent reductions in p300 binding at eNMU which correlated with *NMU* downregulation (Figure 3E). In contrast, e2’s STAT5 mutations only led to a mild decrease in p300 occupancy despite marked reduction in *NMU* expression, suggesting that p300 recruitment is not the primary mechanism for STAT5-mediated facilitator function in eNMU.

### TF-specific regulation of chromatin accessibility and nascent transcription

To gain a deeper understanding of TF-specific regulation of chromatin structure, we performed ATAC-seq on a select set of critical mutants. Focusing on the eNMU locus first, we found distinct changes in chromatin accessibility pattern that seem to be related to the positional context of the disrupted motifs (Figure 4A). Disruption of the GATA1 motif (e1_mGATA1) and the stronger RUNX1 motif (e1_mRUNX1-L)—both situated near the DHS summit—resulted in a broad reduction in ATAC-seq peak height across e1, suggesting that these motifs act as the nucleation sites for chromatin opening. This aligns with the previous reports linking GATA1 and RUNX1 to the recruitment of the SWI/SNF chromatin remodeling complex.^40,41^ Mutating both RUNX1 motifs (e1_mRUNX1) reduced both peak height and peak width at e1, indicating a more severe defect in chromatin decompaction and consistent with its greatest loss in enhancer activity (Figure 3A). In contrast, crippling the more distal RARA/RXRA motifs led to an asymmetrical loss of accessibility on the right flank, while the open chromatin state to their left was likely maintained by GATA1 and RUNX1. Mutations in e2’s STAT5 motif cluster, which resides even further from the DHS center, mainly decreased e2’s accessibility with subtle shrinkage in e1’s peak, suggesting that chromatin opening is not the primary mechanism by which STAT5 facilitates e1’s enhancer activity.

**Figure 4.**
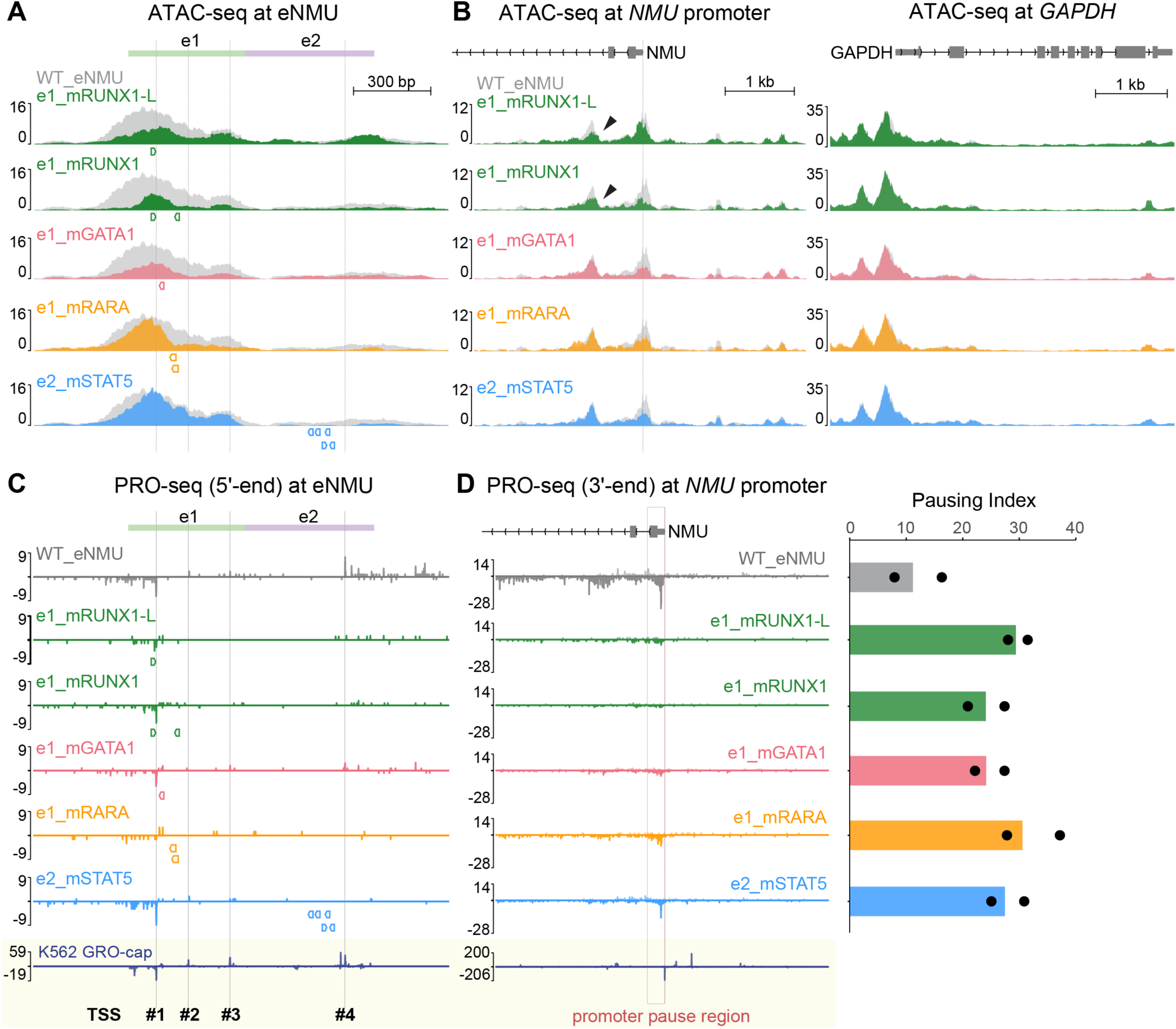
TF-specific regulation of chromatin accessibility and nascent transcription. (A–D) ATAC-seq signal at eNMU (A), *NMU* promoter, and *GAPDH* control locus (B); PRO-seq signal at eNMU (C) and *NMU* promoter (D) in select eNMU mutants. In (B), black arrows highlight the proximal ATAC-seq peak only affected in the RUNX1 motif mutants. In (D), bar = *NMU* pausing index of merged replicates, dots = pausing index of individual replicates. Tracks represent merged biological replicates (n = 2 independent single cell clones). Colored boxes below tracks indicate locations of disrupted TF motifs. Fine vertical lines indicate positions of GRO-cap–defined TSSs (WT K562).^8^ See also Figure S4.

At the *NMU* promoter, all mutants exhibited reduced chromatin accessibility (Figure 4B), consistent with the observed decreases in gene expression. In contrast, the control *GAPDH* gene showed nearly identical accessibility across mutants, demonstrating the reproducibility of our data. Notably, RUNX1 seemed to play an additional role in enhancer–promoter communication, as its motif disruptions specifically affected another *NMU* promoter-proximal ATAC-seq peak (Figure 4B, black arrowheads). Overall, ATAC-seq pattern changes were highly consistent across biological replicates (independent single cell clones) (Figure S4A) and highlight the unique contributions of individual TFs to the chromatin landscape at the enhancer and promoter of *NMU*.

To study TF-specific regulation of nascent transcription at base-pair resolution, we performed PRO-seq on the same set of clones and plotted 5′ positions of PRO-seq reads at the eNMU locus to estimate its TSS usage (Figure 4C). WT_eNMU integration recapitulated the TSS pattern observed in our published K562 GRO-cap data,^8^ validating our methodology. Across the mutants, we found varying degrees of signal reduction, yet the patterns of divergent transcription at e1 and predominantly unidirectional transcription at e2 were largely preserved. Notably, e2’s transcription was diminished not only by its own STAT5 mutations, but also in the e1 mutants where e2’s STAT5 binding was only mildly affected (Figure 3D, right panel). Such decoupling of transcriptional activity from STAT5 occupancy suggests that STAT5 alone is insufficient to drive Pol II initiation at e2. Instead, STAT5 appears to act as an effector that mediates e2’s dependence on e1. Together, these findings depict a highly interconnected transcriptional landscape at the eNMU locus.

Finally, we examined nascent transcription changes at the *NMU* gene by plotting the conventional 3′ ends of PRO-seq reads to represent the locations of paused and elongating Pol II (Figures 4D and S4B). All the mutants showed pronounced signal reductions in both the *NMU* promoter pause region (TSS to TSS+250 bp, Figure 4D, dashed box) and further downstream into the gene body, consistent with their steady-state mRNA levels (Figure 3A). Importantly, the pausing index (PI), defined as the ratio between the pause region and gene body read densities^42^ (Methods), increased 2- to 3-fold in all the mutants compared to WT_eNMU. This points to a defective pause release mechanism at the *NMU* promoter, regardless of specific TF binding at eNMU. Therefore, in addition to its essential role in transcription initiation as demonstrated in the ΔeNMU cell line (Figure 1C), eNMU also regulates Pol II pause release at its target promoter, likely through cofactors shared among its critical TFs.

### Facilitator e2 universally confers enhancer robustness

The extensive crosstalk between e1 and e2 revealed by our functional analysis raised the question on the generality of e2’s facilitator function. To investigate this, we selected eight divergently transcribed, CRISPR-validated distal transcriptional regulatory elements (dTREs) in K562, which showed large effect sizes in the original perturbation studies (Figures S5A and S5B). We assessed their ability to drive *NMU* expression by recombining each dTRE into the eNMU landing pad, either as standalone elements or fused with e2 (Figure 5A). When integrated alone, all the dTREs elevated *NMU* mRNA levels above the baseline of e2 only, although the magnitude of their effects varied drastically (Figure 5B). A closer examination revealed a reasonably good correlation between the dTREs’ intrinsic activities at the eNMU locus and the total number of GRO-cap reads at their endogenous loci (Pearson’s r = 0.85, Figure S5C), suggesting nascent transcription as a reliable indicator of enhancer function. Notably, fusing e2 to the dTREs amplified their activities in every case, with weak elements experiencing greater boosts than strong ones, thereby reducing the variation in their effects. Quantitatively, the intrinsic activities of the dTREs and the amplifications rendered by e2 closely followed a linear log-log distribution (i.e., power-law relationship) (Figure 5C). These observations illustrate a universal buffering function of the facilitator e2 in preventing ultra-low gene expression levels.

**Figure 5.**
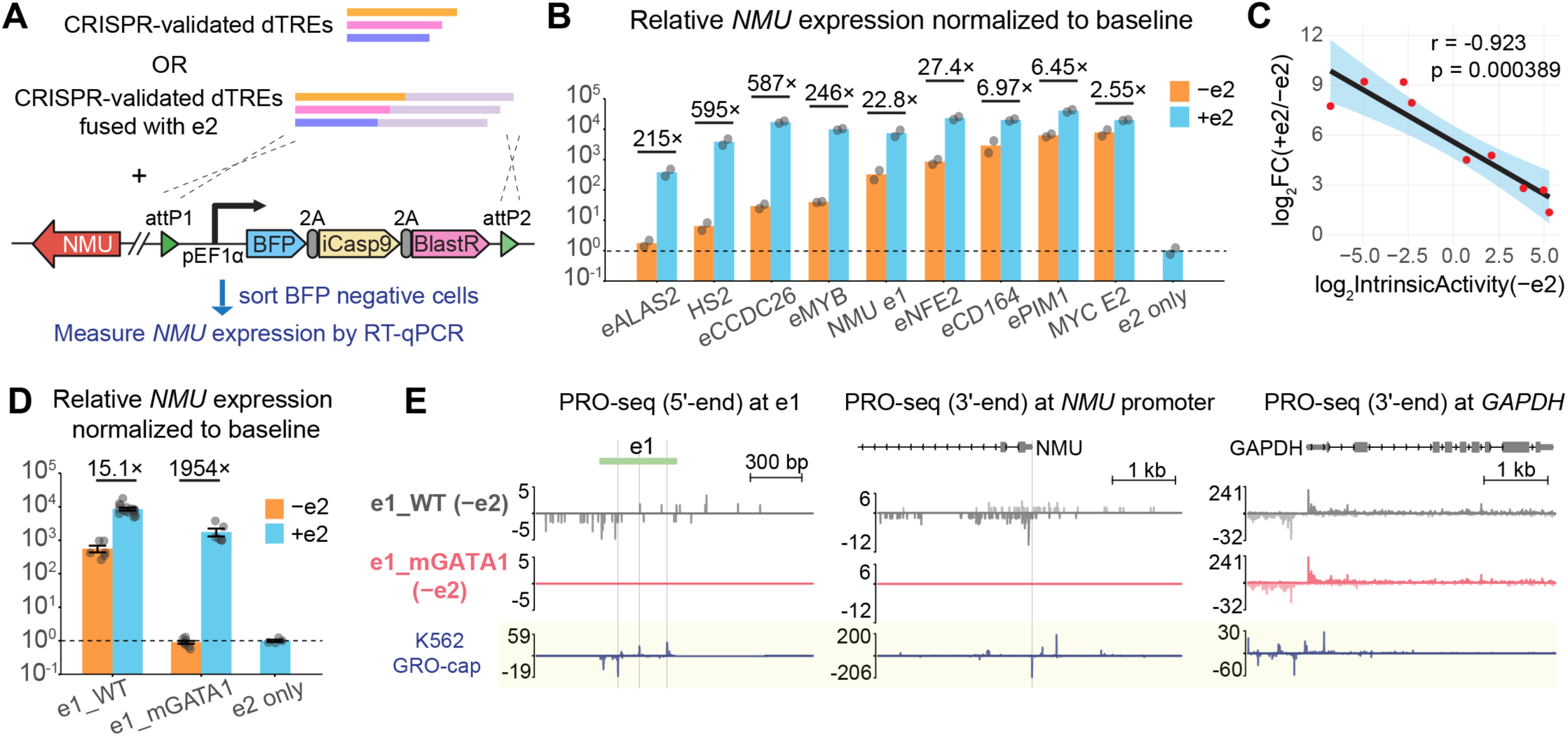
Facilitator e2 universally confers enhancer robustness. (A) Workflow to test e2’s facilitator function on heterologous K562 dTREs using the eNMU landing pad. (B) Enhancer activity of dTREs in the absence or presence of e2, measured by RT-qPCR; e2 only serves as the baseline. n = 2 independent recombination experiments. (C) Correlation between intrinsic activity of elements (−e2) and the fold change with e2 fusion. Pearson’s correlation coefficient (r) and corresponding p-value are shown. Shaded region (blue) indicates the 95% confidence interval around the regression line. (D) *NMU* mRNA levels measured by RT-qPCR in single cell-derived recombinant clones (n ≥ 4) of e1_WT vs. e1_mGATA1 in the absence or presence of e2; e2 only serves as the baseline. Error bars = ± SEM. (E) PRO-seq signal at e1, *NMU* promoter and *GAPDH* control locus in e1_WT and e1_mGATA1 clones lacking e2. Tracks represent merged biological replicates (n = 2 independent single cell clones). Fine vertical lines indicate positions of GRO-cap–defined TSSs (WT K562).^8^ See also Figure S5.

We next asked whether e2’s buffering effect could still apply to a mutated e1 element. Given the critical role of e1’s GATA1 motif in TF cooperativity (Figure 3), we compared the enhancer activities of e1_WT and e1_mGATA1 with or without e2, again in single cell-derived clones. In the absence of e2, disruption of the GATA1 motif completely abolished e1 activity, as reflected by the baseline mRNA level (Figure 5D) and undetectable nascent transcription at both e1 and the *NMU* promoter (Figure 5E). Notably, the presence of e2 restored nascent transcription (Figure 4C and 4D, e1_mGATA1) and rescued the mRNA level by a striking 2,000-fold (Figure 5D), in stark contrast to the 15-fold increase observed for the active enhancer e1_WT. This aligns with the power-law behavior of the heterologous dTREs and highlights the importance of facilitators in safeguarding enhancer robustness against disruptive mutations.

### A 3D regulatory hub of enhancer, promoter and facilitators of *NMU*

In addition to the eNMU region located 94 kb upstream of *NMU*, CRISPRi screens by Gasperini et al.^15^ and Reilly et al.^28^ identified four additional candidate *NMU* “enhancers” at 30.5, 35, 87, and 97.6 kb upstream with varying effect sizes (Figure 6A, purple highlights). We noted that these elements essentially function as facilitators—similar to e2—rather than autonomous enhancers, as they failed to activate *NMU* transcription in the absence of e1 (Figures 1B and 1C, Δe1). In line with this notion, ATAC-seq analysis of the CRISPR deletion lines showed that Δe1, but not Δe2, substantially reduced chromatin accessibility across all the facilitators and the *NMU* promoter to levels comparable to ΔeNMU (Figure 6A, insets). This underscores the hierarchical relationship between the core enhancer e1 and other regulatory elements. Furthermore, public high-resolution intact Hi-C data^3^ shows a distinct stripe pattern anchored at eNMU extending towards the *NMU* promoter (Figure 6B, E–P stripe), suggesting that eNMU actively scans across the 94 kb region and makes widespread contacts. Strong focal interactions, indicated by dot-like patterns, are observed between the *NMU* promoter and F1, eNMU, F3, as well as a distal CTCF/cohesin peak, and also between F1′ and eNMU (Figure 6B, black arrowheads). Consistently, an independent lower-resolution Hi-C study^43^ reveals elevated contact frequencies between almost every pair of the regulatory elements (Figure S6A). These findings together hint at the presence of a spatial regulatory hub for the *NMU* gene (Figure 6D).

**Figure 6.**
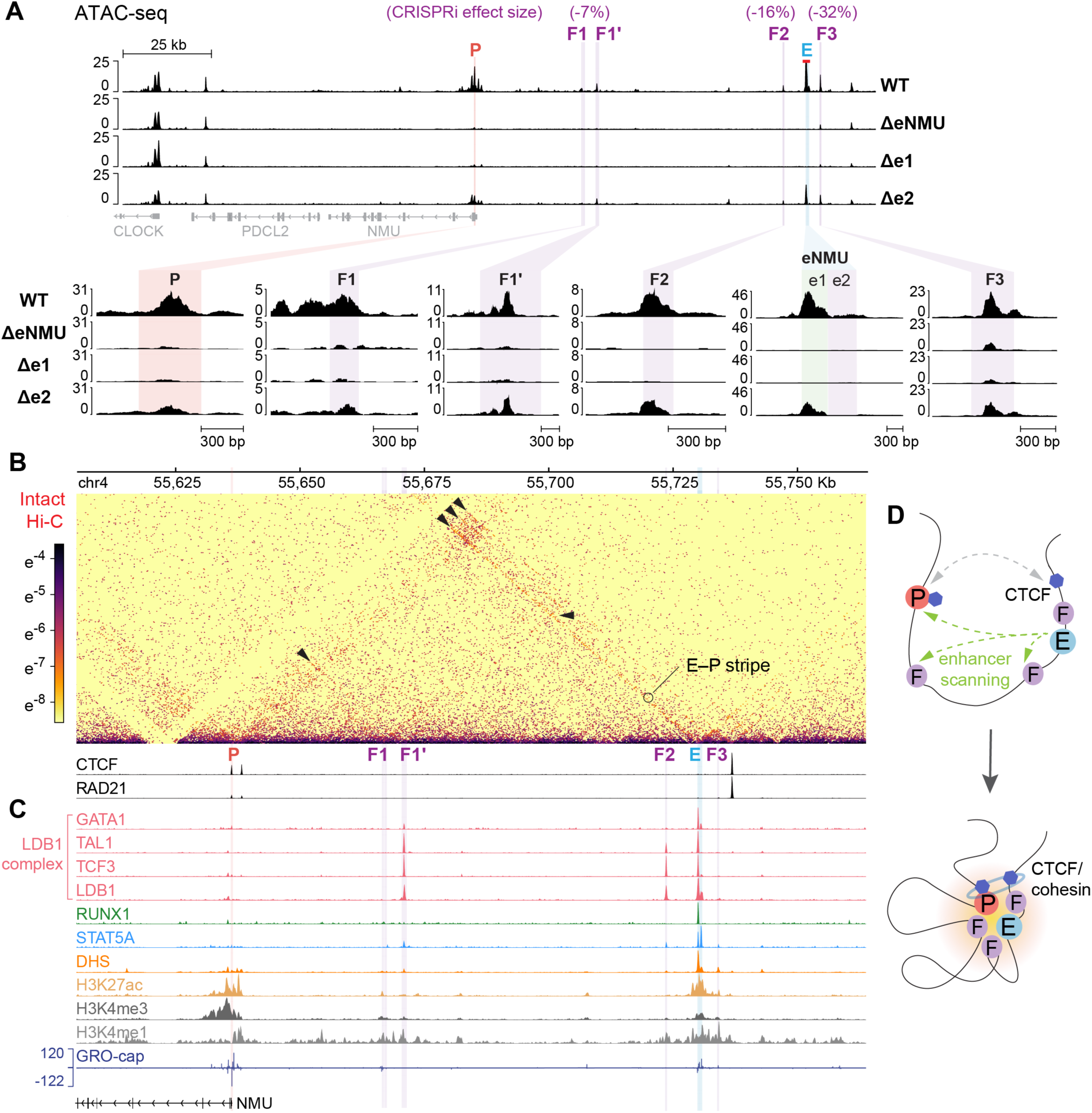
A 3D regulatory hub of enhancer, promoter and facilitators of *NMU*. (A) ATAC-seq signal at the *NMU*–eNMU locus in WT, ΔeNMU, Δe1 and Δe2 cell lines, highlighting *NMU* promoter, eNMU, and facilitators (F1, F2, and F3 from Gasperini et al.^15^; F1 and F1′ from Reilly et al.^28^). Tracks represent merged biological replicates (n = 2 independent cultures). (B–C) Public intact Hi-C (B) and ChIP-seq (B and C)^3,44^ at the *NMU*–eNMU locus in K562. Intact Hi-C is shown at 300-bp resolution, with black arrows indicating pairwise contacts between *NMU* promoter, facilitators, and eNMU; ChIP-seq tracks display signal p-values. Detailed sources and accession information are provided in Table S4. (D) Schematic model illustrating a 3D regulatory hub of enhancer–promoter–facilitator interactions at the *NMU*– eNMU locus. See also Figures S6 and S7.

To explore potential mechanisms underlying the 3D hub formation and facilitator function, we analyzed the epigenomic landscape across the entire *NMU*–eNMU locus, leveraging the vast amount of experimental data available for K562 (Figures 6C and S6B).^3,8,44^ The paucity of the structural proteins CTCF/cohesin at eNMU and its facilitators prompted us to examine the binding of another independent looping factor, the LDB1 complex,^45,46^ at these loci. In erythroid cells, the non-DNA binding transcription cofactor LDB1 forms a stable complex with GATA1, TAL1, E2A/TCF3 transcription factors,^47^ which drives chromatin looping via dimerization of LDB1’s self-association domain.^48^ Indeed, eNMU and two of the facilitators, F1′ and F2, are well occupied by the LDB1 complex (Figures 6C and S6B). Although the *NMU* promoter itself is not bound by LDB1, its downstream proximal DHS peak exhibits low levels of TAL1/TCF3/LDB1 binding (Figure S6B, grey box and inset for *NMU* promoter). Interestingly, instead of GATA1, these factors seem to complex with RUNX1 at this site, which has been reported as an alternative binding partner of TAL1^49^ and LDB1.^50,51^ Of note, we observed decreased accessibility at this promoter-proximal peak exclusively in the e1_mRUNX1 mutants in Figure 4B (black arrowheads), suggesting that RUNX1 binding at eNMU communicates with the RUNX1-containing LDB1 complex near the *NMU* promoter.

Beyond the LDB1 complex, F1′ and F2 also show modest enrichment for STAT5 binding (Figure 6C), which may contribute to their crosstalk with e1, as observed for the facilitator e2. In contrast, the strongest facilitator F3 is predominantly occupied by AP-1 factors along with appreciable binding of the SWI/SNF subunit SMARCA4 (Figure S6B). Nevertheless, signals for other coactivators (p300, BRD4, NCOA1), as well as DHS and H3K27ac, are evidently lower at all four facilitators compared to eNMU. While H3K4me3 is primarily enriched at the *NMU* promoter, all the regulatory loci display comparable levels of H3K4me1, an enhancer mark that has been shown to facilitate enhancer–promoter interactions.^52^ Finally, the minimal GRO-cap signals detected at F1–F3 (Figure 6C), together with the dispensability of TSSs in e2 (Figure 2, Δe2.2, Δe2.4), support the notion that active transcription is not a defining feature of facilitators, thereby solidifying our divergent transcription model for canonical autonomous enhancers. Taken together, our integrative analysis of the constellation of *cis*-regulatory elements at the *NMU*–eNMU locus highlights their spatial connectivity and distinctive epigenomic signatures, providing mechanistic insights into their action.

### Dynamics of eNMU regulation during erythroid differentiation

The remarkable regulatory network of eNMU in K562 led us to explore its function under normal physiological conditions. Given the transcriptomic similarity between K562 cells and early erythroid precursors,^53^ the documented role of NMU peptide in early erythropoiesis,^25^ and the known hematopoietic functions of key TFs acting at eNMU,^37,54–61^ we examined the well-established *ex vivo* erythroid differentiation model of human hematopoietic stem and progenitor cells (HSPCs)^62,63^ (Figure S7A). Reanalysis of published RNA-seq datasets^64–66^ shows that *NMU* is among the most significantly upregulated genes during the differentiation of HSPCs into erythroid precursors (proerythroblasts) (Figure S7B), with expression increasing by over two orders of magnitude (Figures S7C and S7G). This aligns with a recent single-cell multiomics study that reported *NMU* induction during early erythropoiesis of human hematopoietic progenitors.^67^ Importantly, *NMU* induction is accompanied by a progressive increase in the signals of H3K27ac, GATA1, RUNX1,^64^ and chromatin accessibility^66^ at the eNMU locus (Figures S7D and S7F), supporting eNMU as a developmental enhancer of *NMU*. Furthermore, small interfering RNA (siRNA) knockdown of *GATA1* significantly reduces *NMU* expression^64^ (Figure S7E), mirroring the e1_mGATA1 mutant effect in K562 (Figure 3A). Together, these findings highlight the physiological relevance of our results obtained from the immortalized erythroid cell line model K562.

Further examination of the facilitator loci throughout the full differentiation time course^66^ reveals that facilitators F2 and F3 acquire discernible ATAC-seq signals when eNMU accessibility surges (Figure S7F), consistent with their eNMU-dependent behavior in K562 (Figure 6A). Interestingly, the strong facilitator F3 becomes even more accessible during the final stages of erythropoiesis, despite a sharp decline in eNMU accessibility and *NMU* expression. This decoupling of the enhancer–facilitator hierarchy suggests that facilitators work in concert with enhancers in a stage-specific manner and are insufficient to substitute for enhancers in the temporal control of gene expression.

### A putative LTR promoter as a built-in negative regulatory element for enhancer activity

After scrutinizing the activating motifs in eNMU, we shifted our focus to the repressing segment e1.4 (Figure 2) to investigate its functional characteristics. The predominantly unidirectional transcription pattern immediately caught our attention, as opposed to the balanced divergent transcription in the positive regulatory region e1.1–e1.3 (Figure 7A). Interestingly, the entire e1 element corresponds to a MER72 Long Terminal Repeat (LTR) of the ERV1 endogenous retrovirus family. Sequence alignment with the MER72 consensus from the Dfam database^68^ revealed a conserved, weak TATA box variant (CATAA) located 31 bp upstream of the TSS in e1.4, along with a conserved polyadenylation (poly A) signal downstream, matching the typical architecture of an LTR promoter^69^ (Figure 7B). It is thus likely that e1.4 serves as a putative LTR promoter, while e1.1–1.3 functions as its corresponding LTR enhancer. The LTR promoter may compete with the *NMU* promoter for the LTR enhancer activity, which could explain the de-repression of *NMU* gene observed in Δe1.4. To test this hypothesis, we mutated the KLF/SP motifs in either the LTR promoter or the LTR enhancer and assessed their effects in single cell clones (Figure 7C). We chose KLF/SP motifs because (1) disrupting all of them in e1 greatly reduced *NMU* expression in our FlowFISH screen (Figure 2B), and (2) several of them are located closely upstream of TSSs in e1 (Figure 7A), a position generally associated with transcription activation.^70^ Indeed, KLF/SP mutations in the LTR promoter caused a 1.5-fold increase in *NMU* expression (Figure 7C, LTRpr_mKLF), mirroring the effect of Δe1.4. By comparison, KLF/SP mutations in the LTR enhancer or across the entire e1 region dramatically decreased gene expression. We also attempted to strengthen the LTR promoter by optimizing its core promoter elements (CPEs), specifically the TATA box and the Initiator (Inr) motif. As predicted by the competition model, this mutant caused a slight downregulation of *NMU* expression (Figure 7C, mCPE_up). However, attempts to weaken these CPEs showed only a neutral effect, likely due to their inherently weak strength in driving transcription initiation, supporting a dominant role of the KLF/SP motifs in the LTR promoter function.

**Figure 7.**
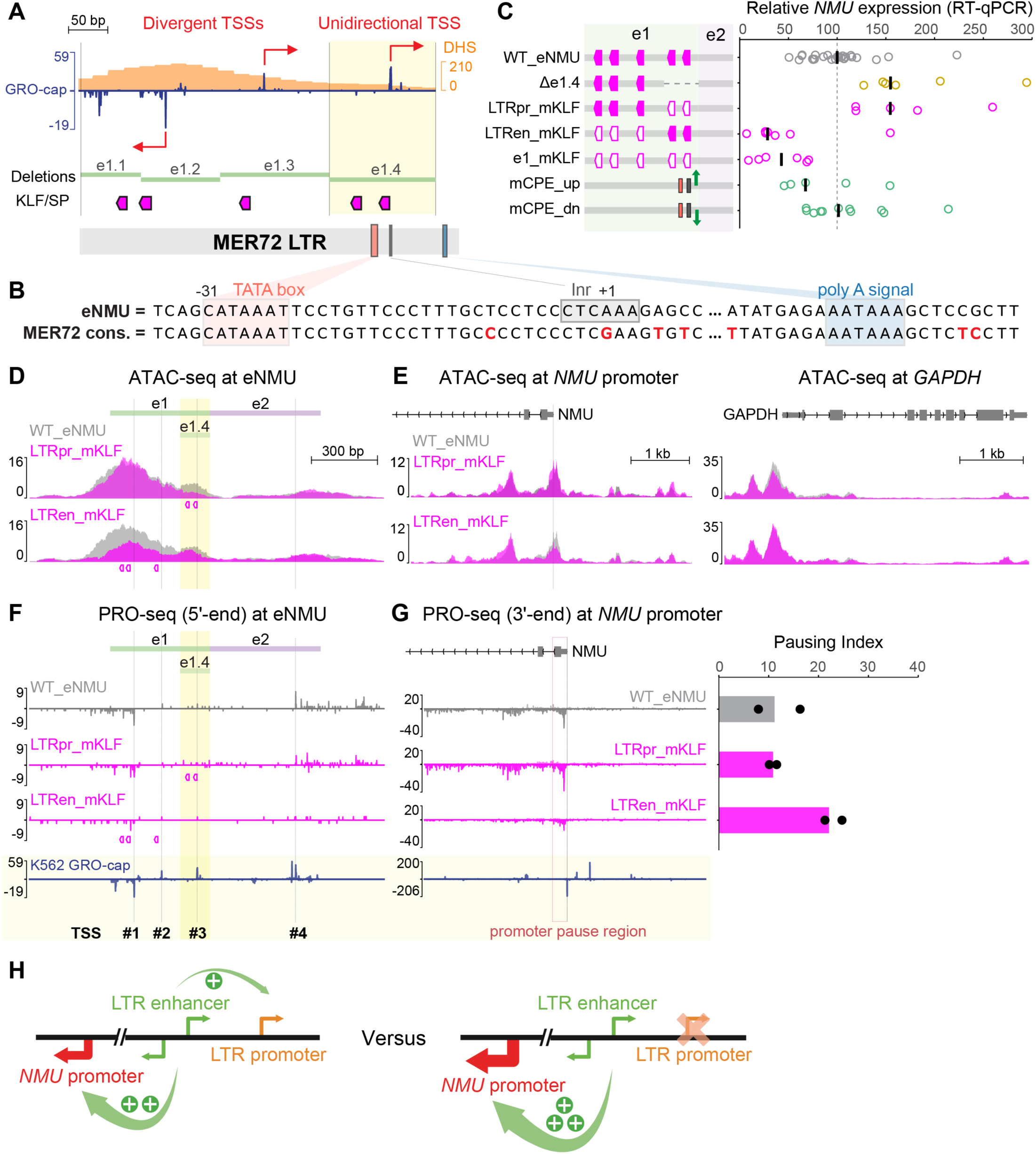
A putative LTR promoter as a built-in negative regulatory element for enhancer activity. (A) Transcription-related sequence features of e1. (B) Sequence alignment between e1 and the MER72 LTR consensus. (C) NMU mRNA levels measured by RT-qPCR in single cell-derived recombinant clones; bars = median. (D–G) ATAC-seq signal at eNMU (D), *NMU* promoter, and *GAPDH* control locus (E); PRO-seq signal at eNMU (F) and *NMU* promoter (G) in select eNMU mutants. In (G), bar = *NMU* pausing index of merged replicates, dots = pausing index of individual replicates. Tracks represent merged biological replicates (n = 2 independent single cell clones). Colored boxes below tracks indicate locations of disrupted TF motifs. Fine vertical lines indicate positions of GRO-cap–defined TSSs (WT K562).^8^ (H) Proposed competition model between the LTR promoter and the *NMU* promoter. See also Figure S4 and Table S1.

To exclude the possibility that the KLF/SP sites in e1.4 act as repressor motifs, we performed ATAC-seq in both the LTR promoter and the LTR enhancer mutants. This revealed highly localized accessibility reductions confined to the respective mutated regions (Figures 7D and S4A), confirming the activating function of KLF/SP in both contexts. However, accessibility at the *NMU* promoter changed in opposite directions (Figure 7E), supporting the idea that the putative LTR promoter and enhancer operate as distinct regulatory elements for *NMU*, likely through a promoter competition mechanism (Figure 7H).

Finally, we examined the nascent transcription profiles of the KLF/SP mutants by PRO-seq. At the eNMU locus, the LTR promoter mutant showed a nearly identical pattern to WT_eNMU, suggesting unperturbed Pol II recruitment (Figure 7F). Furthermore, the pausing index of *NMU* also remained unaltered due to the proportional increase in the pause region and gene body reads (Figures 7G and S4B). Conversely, the LTR enhancer mutant resembled other e1 mutants studied in Figure 4, exhibiting reduced eNMU and *NMU* signals, along with a doubled pausing index. These observations thereby raise an interesting possibility: promoter competition could provide a unique advantage in modulating target gene transcription while maintaining normal Pol II pause–release dynamics.

## Discussion

In this study, we established a novel landing pad platform to systematically interrogate the molecular architecture of the potent long-range enhancer eNMU at its native locus. Through detailed functional dissections of key mutants and integrative mining of public datasets, we uncovered several recurring themes supported by multiple lines of evidence: (1) TFs exert unique and cooperative functions in a context-specific manner; (2) facilitators depend on the core enhancers while ensuring robustness of their enhancer partners; (3) divergent transcription accurately demarcates active enhancer units. Collectively, our findings illuminate an intricate and coordinated interplay among distinct classes of *cis*-regulatory elements—enhancers, facilitators, and promoters—that underpins precise transcriptional regulation.

Previous enhancer studies have primarily employed approaches such as random mutagenesis,^11,12,32,71^ tiling disruptions,^33,72,73^ and specific motif manipulations^74–77^ to identify functional features within regulatory elements. In contrast, our study applied a distinct framework to dissect eNMU, grounded in our divergent transcription-based unit definition of active human enhancers.^8^ Building on the foundational work of Gasperini et al.^15^ and Tippens and Liang et al.,^9^ this approach enabled us to progressively refine the bona fide *NMU* enhancer unit from the full eNMU region to its sub-element e1, and ultimately to a minimal, divergently transcribed LTR enhancer core (Figure 7). Importantly, the core enhancer activity is modulated by the surrounding sequence features within eNMU—specifically, augmented by the intrinsically inactive facilitator element e2 and repressed by the adjacent unidirectionally transcribed LTR promoter. These findings reveal the regulatory complexity of the eNMU locus and highlight the strength of our divergent transcription model in precisely delineating functional enhancer units, thereby guiding the future classification of diverse distal regulatory elements.

Unlike previously described facilitators identified in the context of hyper-active super-enhancers,^23,24^ the presence of multiple facilitators associated with a typical enhancer across the ∼100-kb *NMU*–eNMU region raises the possibility that many CRISPRi-identified “enhancers” may in fact function as facilitators. Notably, the eNMU-associated facilitators exhibited virtually no intrinsic activity even when present all together in the genome (0.01% of WT expression, Figure 1B, Δe1), consistent with their inherently weak, enhancer-dependent accessibility patterns (Figure 6A). This stands in contrast to the facilitators within super-enhancers, which tend to display strong signals of open chromatin, TF/coactivator binding, and Pol II recruitment—features likely contributing to their residual intrinsic enhancer activity.^24^ We speculate that a continuum of enhancer potential exists along the enhancer– facilitator spectrum, with the eNMU-associated facilitators situated at the extreme low-activity end. The lack of intrinsic activity is crucial in confining facilitator function to potentiating and buffering pre-established enhancers (Figure 5) while preventing ectopic gene activation. With the continuing advances in genome engineering technologies, it will be increasingly important to interrogate all candidate regulatory elements simultaneously within their native chromatin hub environments, to distinguish autonomous enhancers from affiliated facilitators and to better understand their mechanistic interplay.

In addition to the enhancer–facilitator axis, the LTR enhancer–promoter axis within eNMU represents another potentially widespread and underappreciated mode of *cis*-regulatory behavior, especially considering the high abundance of LTRs in the human genome and their nearly 10% representation of all ENCODE candidate *cis*-regulatory elements (cCREs).^78^ By leaving the enhancer intact, the LTR promoter can compete with the gene promoter without disrupting normal Pol II pause–release dynamics, instead simply siphoning transcriptional activity toward itself. This regulatory strategy enables fine-tuning of target gene expression, particularly when the LTR promoter harbors motifs for developmental stage-specific TFs. Moving forward, a genome-wide search for functional unidirectional TSSs, including but not limited to those derived from LTR promoters, will be crucial for constructing a more comprehensive map of transcriptional networks.

How do these functionally distinct classes of *cis*-regulatory elements coordinate to achieve precise and robust transcriptional regulation? We propose that the answer lies in the combinatorial action and synergy of a repertoire of *trans*-acting TFs and cofactors. At the core LTR enhancer unit, we observed strong cooperative binding of key TFs GATA1 and RUNX1 (Figures 3B and 3C), despite their motifs being separated by ∼40 bp—a spacing that likely limits direct protein–protein interactions. This suggests that indirect mechanisms^79,80^ may underlie their cooperation, including DNA conformational changes,^81^ co-binding to a shared cofactor or a multiprotein complex,^79^ and nucleosome-mediated collaborative competition.^82–85^ At the adjacent LTR promoter, activating KLF/SP family TFs played a context-specific role to compete for enhancer activity (Figure 7). At the facilitator e2, STAT5 binding to a tandem array of five motifs critically amplified e1’s enhancer activity, despite modest impact on e1’s chromatin accessibility, transcription initiation, and p300 recruitment (Figures 3 and 4). It is worth noting that e2’s own accessibility and transcription depended not only on its own STAT5 binding, but also on the integrity of e1’s key TF binding (Figures 4A and 4C). This peculiar behavior of STAT5 echoes the recently proposed concept of “context-only” TFs,^86^ which do not provide DNA access themselves but instead amplify the activity of “context-initiator” TFs by establishing cooperative environments. These two classes of TFs partner promiscuously without requiring close motif proximity, consistent with e2’s universal buffering effect on various heterologous enhancers (Figure 5). We speculate that STAT5 binding at e2 fosters multivalent interactions^87,88^ with e1-bound TFs via its intrinsically disordered C-terminal transactivation domain,^89^ thereby enhancing the “stickiness” of the regulatory hub. Nonetheless, we cannot exclude the possibility that STAT5 engages some unique coactivators which have yet to be identified.

In addition to the disordered C-terminal transactivation domain, STAT5 contains an N-terminal oligomerization domain that allows tetramerization of active STAT5 dimers on tandemly linked motifs.^90,91^ This oligomerization extends STAT5’s DNA binding specificity to low-affinity sites,^92^ which may explain the ∼30% residual binding observed upon mutating two conserved bases in all five STAT5 motifs at e2 (Figure 3D, right panel). Such oligomerization might also facilitate spatial connectivity between eNMU and the STAT5-bound facilitators F1′ and F2—reminiscent of GAGA-associated factor (GAF) oligomerization at a subset of tethering elements in developing *Drosophila* embryos,^93^ which, despite lacking intrinsic enhancer activity, are essential for long-range enhancer–promoter communication.^94^ In parallel, dimerization of the LDB1 complex bound at F1′ and F2, the enhancer e1, and the *NMU* promoter (Figures 6 and S6) may further promote chromatin contacts between these loci. Together, these potential mechanisms suggest a broader architectural role for facilitators in organizing 3D regulatory hubs independent of CTCF or cohesion, underscoring an exciting avenue for future investigation.

A final noteworthy observation from our functional analysis is that, despite substantial variation in chromatin accessibility pattern at eNMU across different motif mutants (Figure 4A), the transcriptional output and Pol II pause–release dynamics (pausing index) of *NMU* were altered to similar extents (Figures 3A and 4D). Such decoupling between enhancer accessibility and gene activation is consistent with prior findings^82,95^ and highlights the importance of specific TF inputs and their associated cofactors in driving functionally productive enhancer–promoter communication. Future work should aim to define the full repertoire of these regulatory components and the steps of transcription that they influence (such as chromatin opening, Pol II initiation, and pause release).

In summary, we conducted a rigorous *in situ* dissection of a robust long-range enhancer at unprecedented architectural resolution, providing experimental evidence that resonates with and extends current models of enhancer function. The intricate crosstalk among spatially and functionally linked *cis*-regulatory elements—including enhancers, facilitators, and promoters—underscores the importance of a holistic framework to decode their mechanistic interplay. We anticipate that our efficient and versatile recombinase-mediated genome rewriting platform will serve as a powerful tool to drive these efforts forward.

### Limitations of the study

Our motif mutagenesis approach could not definitively identify the functional TFs acting at eNMU. For instance, both GATA1 and GATA2 are well expressed in K562 cells and bind similar/identical motifs, making it difficult to distinguish their individual contributions. We attributed the observed effects to GATA1 in our study, because it is the most highly expressed GATA family factor in K562^96^ and the master regulator of erythropoiesis. Similarly, we did not determine the exact TFs binding the critical RARA/RXRA motifs, given the low abundance of RARA and RXRA proteins^96,97^ and the presumed absence of retinoic acid signaling in K562 under standard culture conditions. It is possible that other nuclear receptors recognize and bind these motifs. In addition, our mutagenesis screen may have missed some functional motifs, as it relied on the availability of ChIP-seq data to confirm TF binding. Furthermore, while we made every effort to avoid disrupting overlapping motifs, some degree of interference was unavoidable—for example, between the right RUNX1 motif and the adjacent RARA/RXRA motif. As we introduced only a single version of the transversion mutations, we also cannot completely exclude the possibility of inadvertently creating novel TF binding sites, despite efforts to minimize matches to known motifs. Finally, although e2 displayed power-law buffering behavior across eight heterologous enhancers, larger-scale studies are warranted to fully capture the complexity of enhancer–facilitator synergism.

## RESOURCE AVAILABILITY

### Lead contact

Further information and requests for resources and reagents should be directed to and will be fulfilled by the lead contact, John T. Lis.

### Materials availability

Plasmids and cell lines generated in this study are available upon request.

### Data and code availability

- All raw and processed next-generation sequencing data have been deposited in the Gene Expression Omnibus (GEO) under accession numbers GSE299351, GSE299353, GSE299354, GSE299355.
- This paper does not report original code.
- Any additional information required to reanalyze the data reported in this paper is available from the lead contact upon request.

## ACKNOWLEDGMENTS

We thank all past and present members of the Lis Lab for insightful discussions and support throughout this work. A special thank you to Dr. Judhajeet Ray, former research associate in the Lis Lab and currently at the Broad Institute, for his unwavering mentorship and patience in guiding Zhou Zhou during the formative early years of her Ph.D. training. Another special thank you to Jessica West, former Ph.D. candidate in Dr. Andrew Grimson’s lab and currently at UCSF, for generously sharing landing pad-related plasmids and for valuable input on the design of the recombination platform and library screening strategy. Thanks to former lab members Dr. Jin Liang and Nathaniel Tippens in Dr. Haiyuan Yu’s lab for providing eNMU deletion cell lines and contributing early ideas to the construction of the landing pad system. Thanks to Dr. Andrew Grimson and Dr. Charles Danko for their critical feedback on the manuscript. Thanks to Adam He (Dr. Charles Danko’s lab) and Haining Chen (Dr. Franklin Pugh’s lab) for helpful discussions and expertise. Thanks to Xinchen Chen (Dr. Chun Han’s lab) for guidance on figure preparation using Adobe Illustrator. Fluorescence-activated cell sorting experiments were conducted at the Cornell Institute of Biotechnology’s Flow Cytometry Facility. Most next-generation sequencing data were generated by the Institute’s Epigenomics Core Facility, and initial pilot data produced by the Institute’s Genomics Facility. This work was supported by the National Human Genome Research Institute (NHGRI) grant 5R01HG012970 to J.T.L. and H.Y.

## AUTHOR CONTRIBUTIONS

Z.Z., J.T.L., and A.O. conceptualized the study. Z.Z. designed the eNMU mutagenesis scheme with substantial input from J.T.L., A.O., J.Z., and H.Y. Z.Z. and A.L. constructed the eNMU mutant library. Z.Z. generated the landing pad cell lines, performed all functional genomics experiments with assistance from A.O., and conducted all data analyses. Z.Z., J.T.L., and A.O. interpreted the data and wrote the manuscript, with all authors contributing to revisions. J.T.L. and H.Y. secured funding for the study.

---

**Figure S1.**
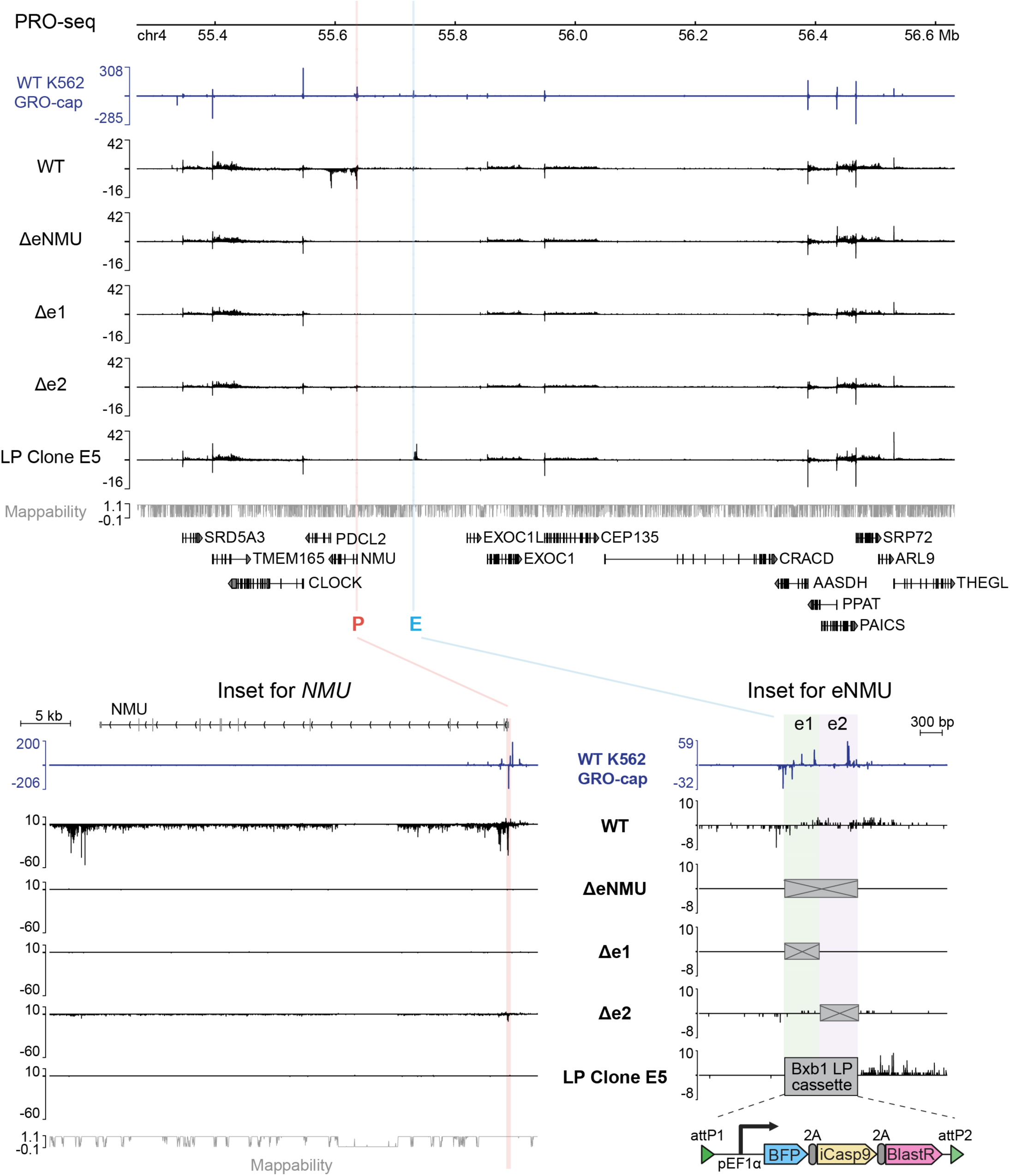
eNMU specifically regulates *NMU* gene transcription in K562. PRO-seq (3′-end) tracks of WT, ΔeNMU, Δe1, Δe2 and eNMU LP cell lines across a ∼1 Mb region around *NMU*; insets show the full *NMU* gene and eNMU loci. Highlighted regions indicate *NMU* promoter (P) and eNMU (E). Note that at the eNMU locus in LP Clone E5, the prominent PRO-seq signal downstream (to the right) of the Bxb1 LP cassette originates from readthrough transcription driven by the strong EF1ɑ promoter within the selection cassette. However, transcriptional activity of the LP did not exhibit any enhancer function to activate the distal *NMU* gene. Tracks represent merged biological replicates (n = 2 independent cultures). WT K562 GRO-cap data^8^ shown as the TSS reference. Related to Figure 1.

**Figure S2.**
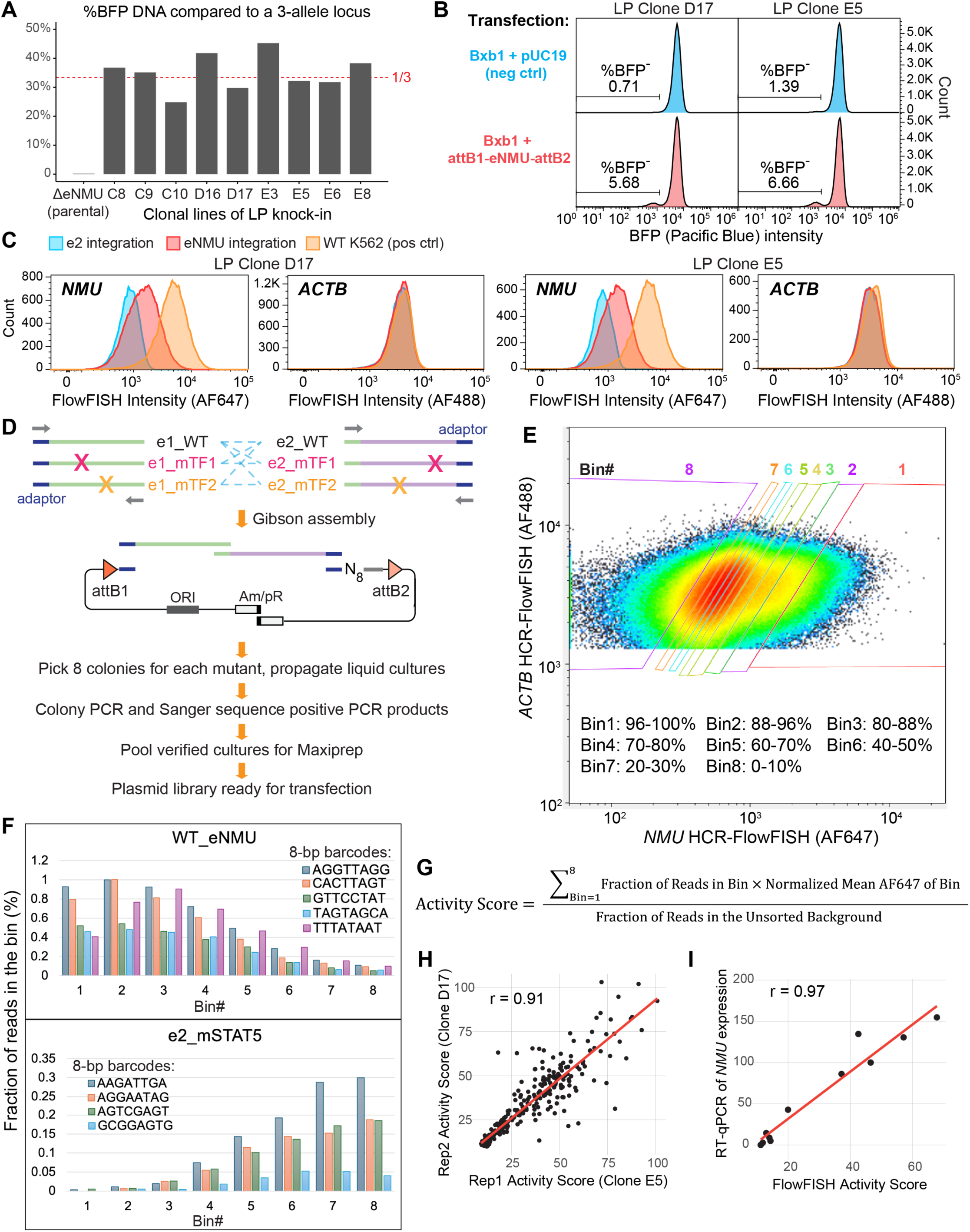
eNMU landing pad construction and mutant screen. (A) LP copy number analysis by quantitative PCR on genomic DNA (BFP DNA vs. 3-allele control locus). (B) Bxb1 recombination efficiency measured by BFP loss in two independent LP clones using flow cytometry. (C) Validation of HCR-FISH in the same LP clones as in (B); *ACTB* served as the housekeeping gene control. (D) Gibson assembly workflow for constructing the barcoded eNMU mutant library. (E) Flow cytometry binning strategy using *ACTB* as an internal control for cell size, transcription level, and staining efficiency. (F) Barcode distribution across 8 sorting bins for two example elements in the mutant library: WT_eNMU and e2_mSTAT5. (G) Calculation of activity scores using a weighted average of barcode distributions. (H) Correlation of median barcode activity scores between biological replicates (n = 2 independent LP clones subjected to recombination and FlowFISH). Pearson’s correlation coefficient (r) is shown. (I) Correlation between FlowFISH-measured activity scores and RT-qPCR quantifications of select mutants, based on median activity values from each assay. Pearson’s correlation coefficient (r) is shown. Related to Figures 1 and 2. See also Methods.

**Figure S3.**
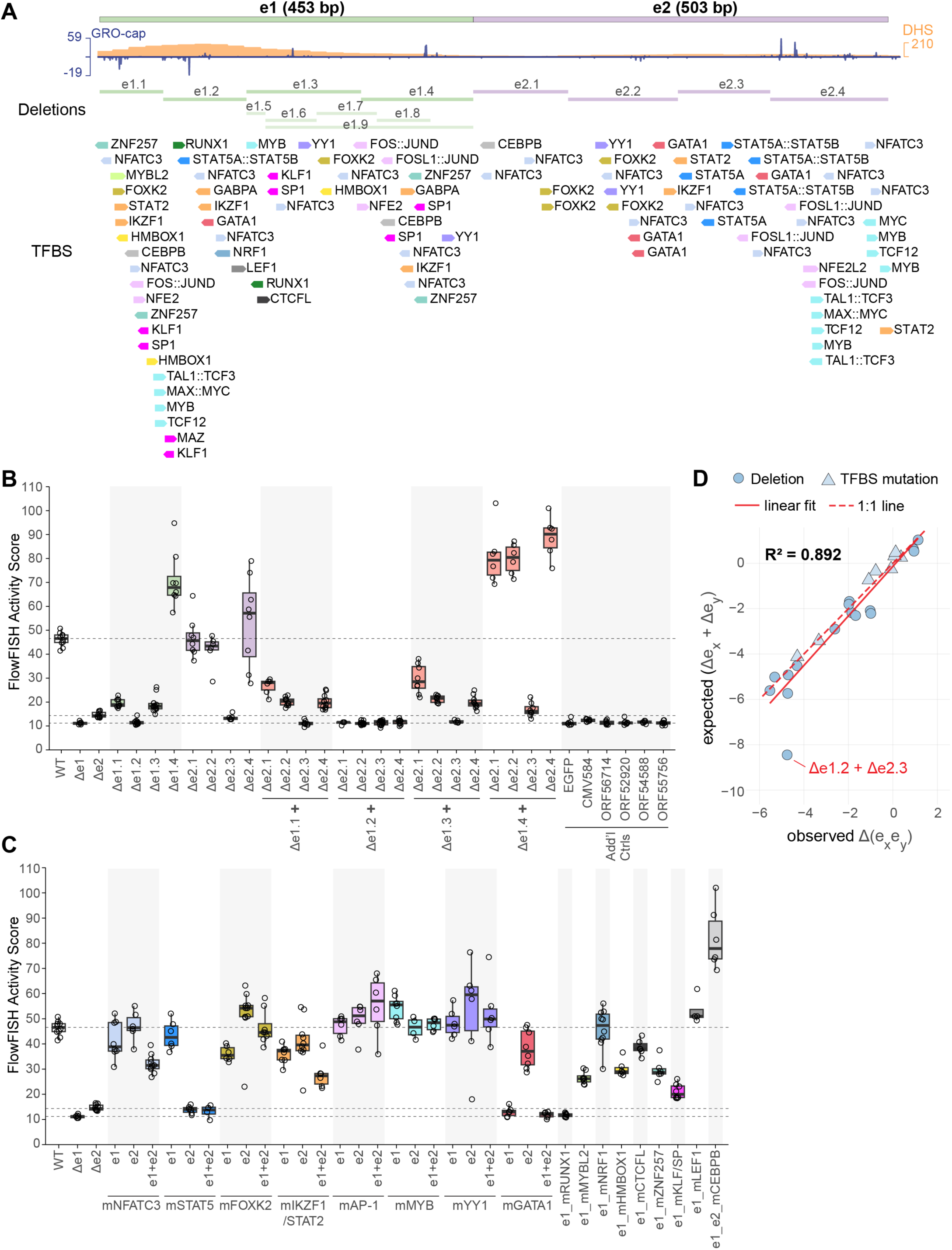
Complete eNMU mutant screen results reveal a multiplicative model of double mutants’ effects. (A) Full eNMU mutant design separating overlapping motifs of different TFs. (B) FlowFISH-measured activity scores of single and double deletions, together with additional exogenous control elements. (C) FlowFISH-measured activity scores of all TF motif mutations in e1, e2, or both. Note that the e1_e2_mCEBPB mutant exhibited higher enhancer activity than WT_eNMU, which may be attributed to an LTR promoter-mediated mechanism as illustrated in Figure 7. (D) Linear regression of observed log_2_ fold changes in double mutants vs. expected additive effects (sum of log_2_ fold changes from corresponding single mutants). R² from linear regression is shown. Dashed 1:1 line indicates perfect additivity. Highlighted outlier: expected effect of the Δe1.2+Δe2.3 mutant fell below FlowFISH’s detection range, preventing assessment of additivity. Related to Figure 2. See also Methods, Tables S1 and S3.

**Figure S4.**
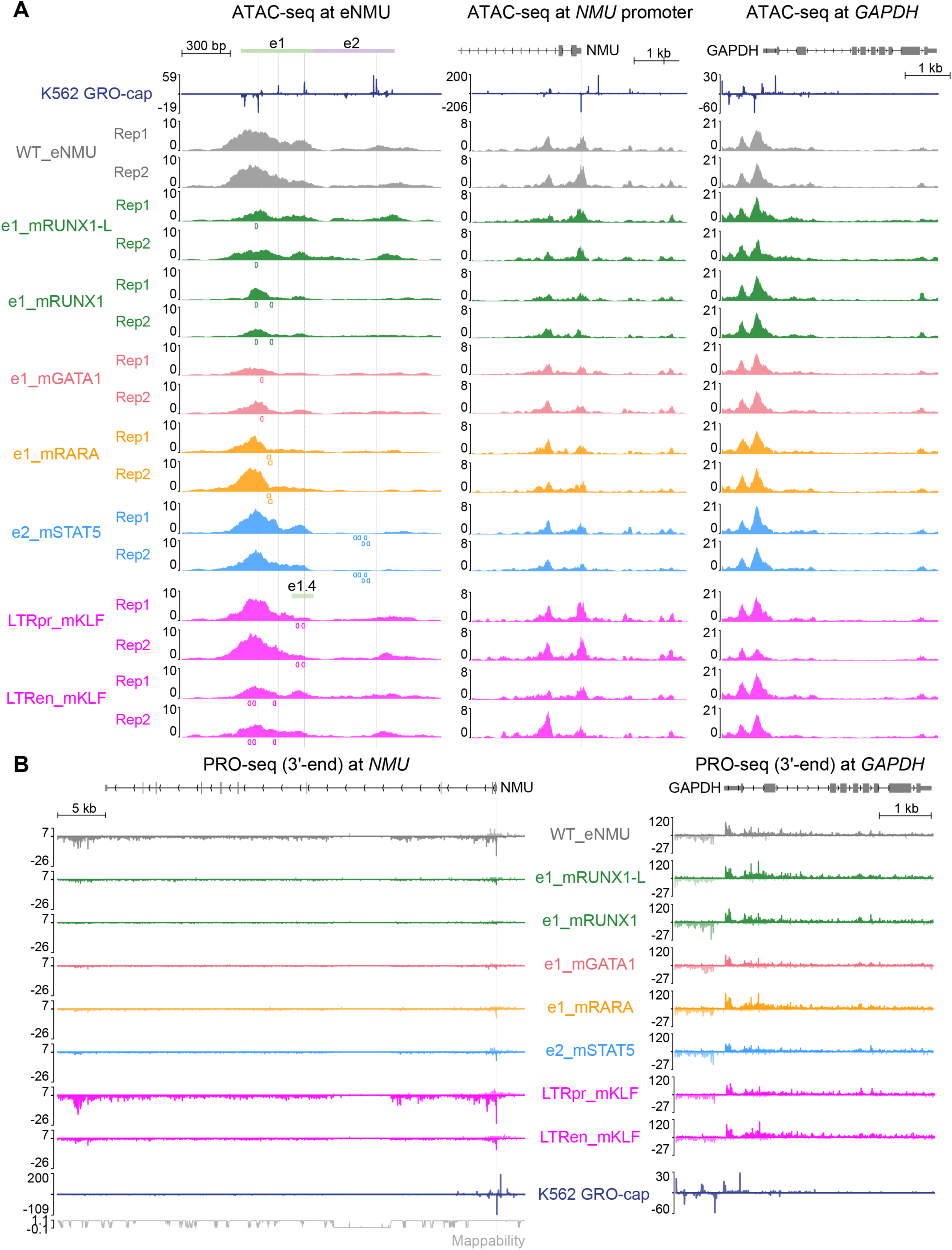
Reproducibility of ATAC-seq and PRO-seq data across clones and mutants. (A) ATAC-seq signal at eNMU, *NMU* promoter and *GAPDH* control locus for two independent single cell-derived clones of all analyzed eNMU mutants in this study. Colored boxes below tracks indicate locations of disrupted TF motifs. (B) PRO-seq tracks at the full *NMU* and *GAPDH* (control) genes in the same mutants as in (A). Tracks represent merged biological replicates (n = 2 independent single cell clones). Fine vertical lines indicate positions of GRO-cap–defined TSSs (WT K562).^8^ Related to Figures 4 and 7.

**Figure S5.**
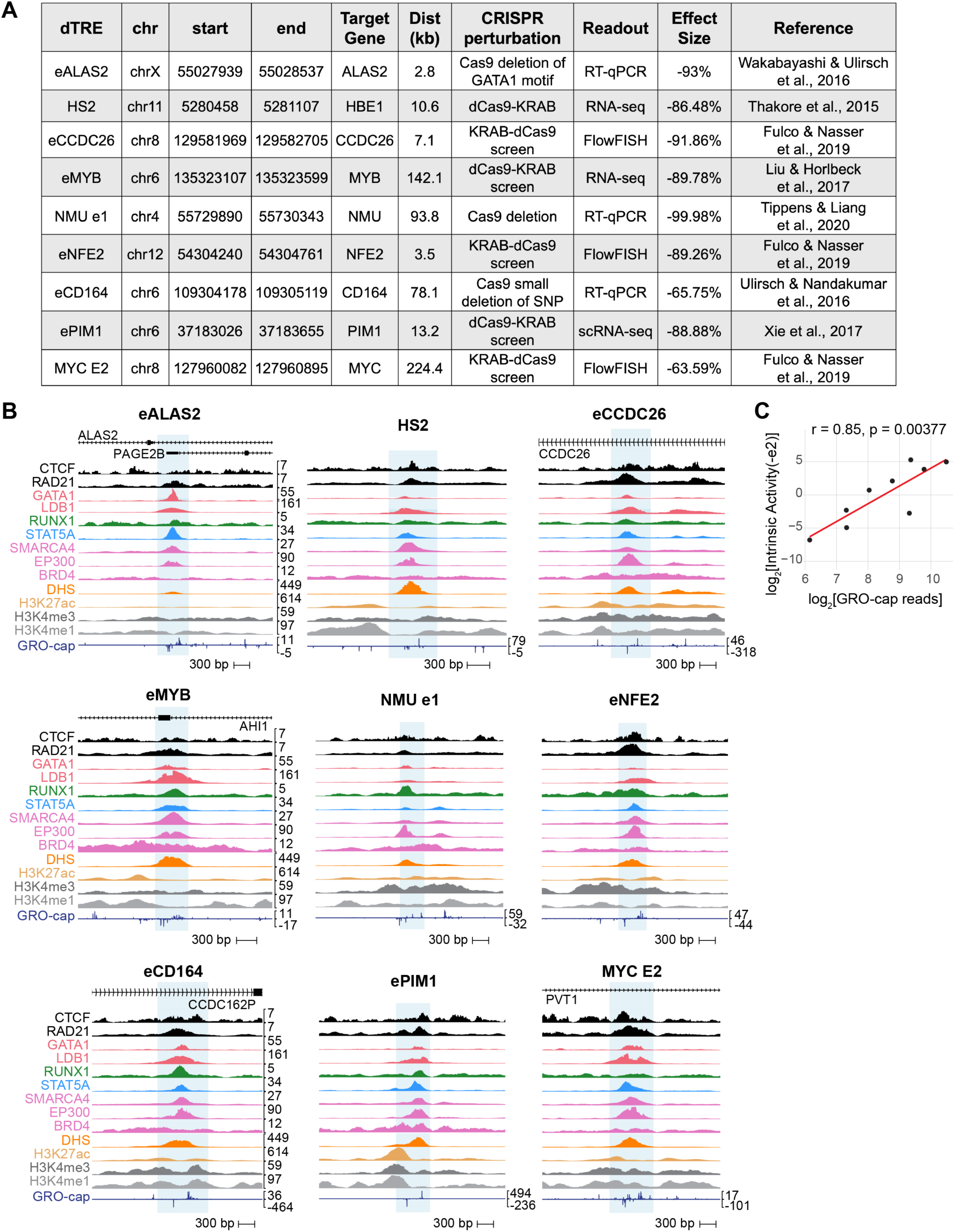
Testing CRISPR-validated heterologous K562 dTREs at the eNMU locus. (A) Summarized information of selected K562 dTREs from previous studies. Effect sizes obtained from Fulco et al.^16^ (except for NMU e1, which is from Tippens and Liang et al.^9^). (B) Native genomic contexts of each dTRE; tested regions highlighted in light blue. Track scales are consistent across dTRE regions, except for GRO-cap,^8^ which uses an individually indicated scale. Detailed sources and accession information are provided in Table S4. (C) Correlation between GRO-cap read counts at dTREs and their intrinsic enhancer activity (−e2) at the eNMU locus. Pearson’s correlation coefficient (r) and corresponding p-value are shown. Related to Figure 5.

**Figure S6.**
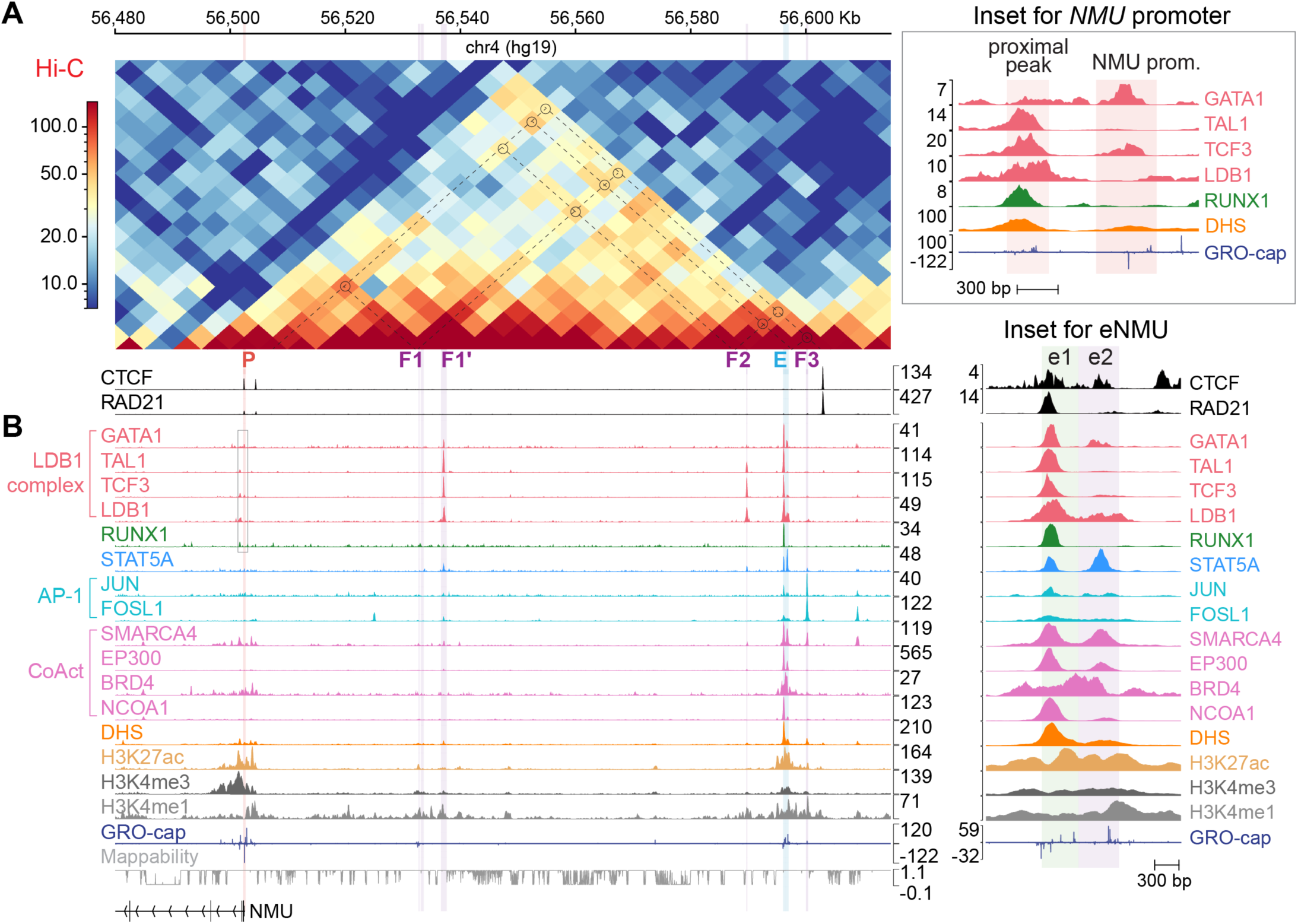
Additional epigenomic features of enhancer, promoter and facilitators. (A) Public Hi-C^43^ and ChIP-seq tracks^3^ of CTCF and RAD21 at the *NMU*–eNMU locus in K562. Hi-C is shown at 5-kb resolution, with dashed lines and open circles marking pairwise contacts between *NMU* promoter, facilitators, and eNMU. Note that some contact anchors may not align perfectly with the regulatory elements, possibly due to the limited resolution of this Hi-C dataset. (B) Expanded ChIP-seq tracks^3,44^ displaying signal p-values. Grey box highlights the *NMU* promoter and its proximal region, with a zoomed-in view shown on the top right. A separate inset on the right zooms in at the eNMU region, shown at the same scale as the full locus, except where otherwise indicated. Detailed sources and accession information are provided in Table S4. Related to Figure 6.

**Figure S7.**
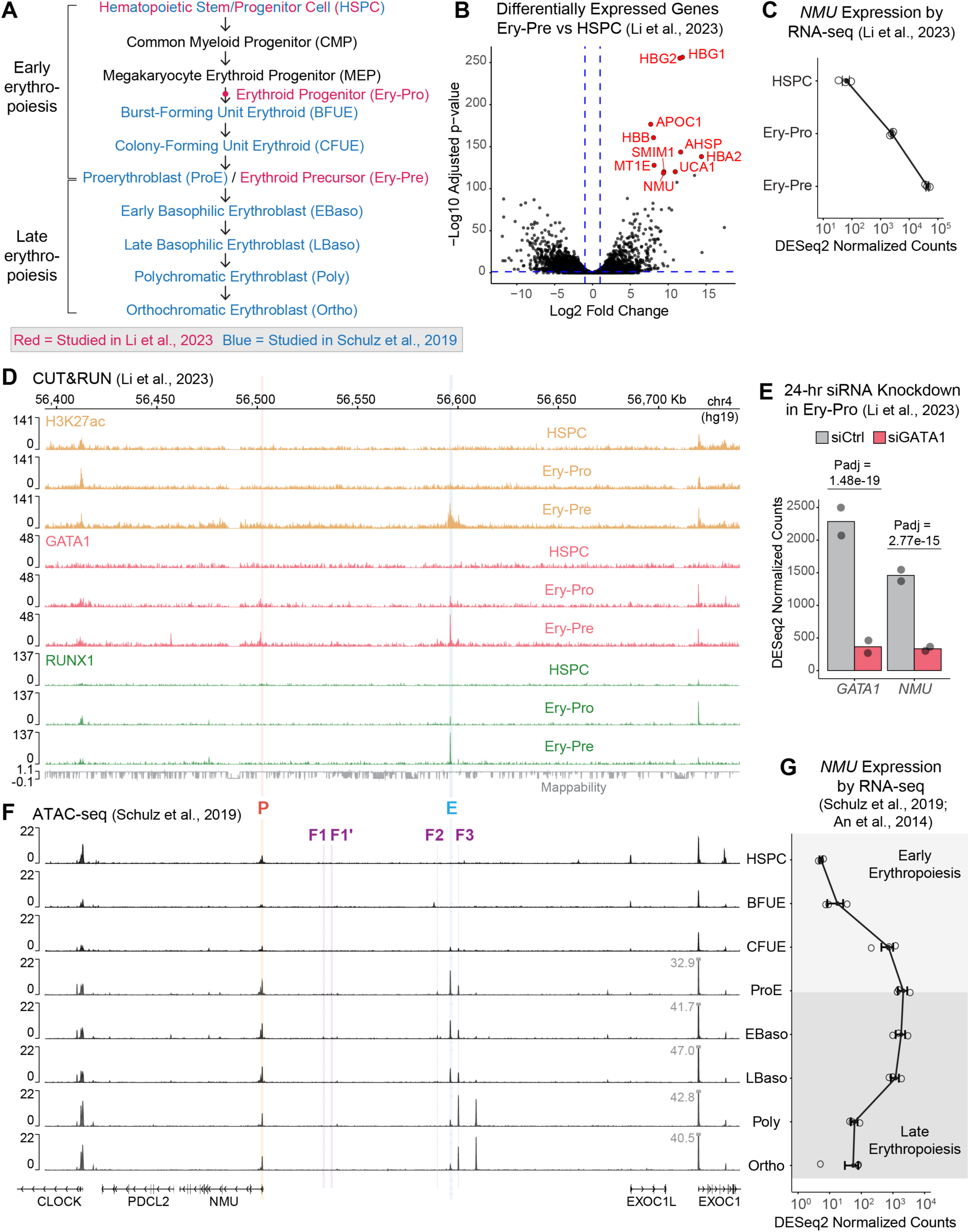
Dynamics of eNMU regulation during erythroid differentiation. (A) Stages of HSPC erythroid differentiation analyzed in Li et al.^64^ (red) and Schulz et al.^66^ (blue). (B) Volcano plot showing genome-wide expression changes between HSPC and Ery-Pre stages, reanalyzed from Li et al.^64^ RNA-seq data. Horizontal and vertical blue lines mark adjusted p = 0.05 and log₂ fold changes of ±1, respectively. Red dots highlight the top 10 most significantly upregulated genes, including the key erythroid markers *HBG1* and *HBG2* (β-like globin genes). (C) *NMU* expression changes during early erythropoiesis, reanalyzed from Li et al.^64^ RNA-seq data (n = 3). (D) CUT&RUN signal of H3K27ac, GATA1 and RUNX1 at the *NMU*–eNMU locus during early erythropoiesis. Tracks show one representative biological replicate from Li et al.^64^ (E) *GATA1* and *NMU* expression changes following 24-hr siRNA knockdown of *GATA1* in Ery-Pro cells, reanalyzed from Li et al.^64^ RNA-seq data (n = 2). (F) ATAC-seq signal at the same locus as in (D) throughout the full HSPC differentiation time course. Tracks show merged biological replicates (n = 2) from Schulz et al.^66^ (G) *NMU* expression changes during the same stages as in (F), reanalyzed from An et al.^65^ and Schulz et al.^66^ RNA-seq data (n = 3). Related to Figure 6.

## EXPERIMENTAL MODEL AND STUDY PARTICIPANT DETAILS

### Cell lines and culture

Parental wildtype K562 cells, an immortalized erythroleukemia cell line isolated from the bone marrow of a 53-year-old female patient with chronic myelogenous leukemia (CML), were obtained from the America Type Culture Collection (ATCC) (ATCC Number CCL-243) by the Yu lab and generously provided to us. Genetically modified, homozygous eNMU deletion lines (ΔeNMU, Δe1, and Δe2) were also kind gifts of the Yu lab. All the other engineered K562 cell lines, including the eNMU landing pad lines and single cell-derived recombinant clones, were generated by this study (see METHOD DETAILS). All the K562 lines were cultured in RPMI 1640 media supplemented with GlutaMAX (Gibco) and 10% heat-inactivated FBS (Avantor) at 37°C with 5% CO_2_ in a humidified sterile incubator. Cell density was maintained between 0.1 ∼ 1 × 10⁶ cells/mL, and mycoplasma testing was performed routinely.

### Transfection and cell sorting

All transfection experiments in K562 cells were carried out using Lonza’s Nucleofector 2b device and the Nucleofection Kit V, following manufacturer’s instructions. Specifically, one single cuvette was used to transfect 1 million cells with a total of 5 μg plasmid DNA; for co-transfection of two plasmids, 2.5 μg of each plasmid was used. All cell sorting experiments were performed on the Sony MA900 Multi-Application Cell Sorter using a 100-μM chip (catalog no. LE-C3210; Sony).

## METHOD DETAILS

### Single-copy eNMU landing pad cell line construction

To CRISPR knock in the Bxb1 landing pad at the eNMU locus in an eNMU-null background, we first amplified the genomic region surrounding the eNMU locus in the ΔeNMU cell line, inserted it into a HindIII-linearized pEGFP-N1 vector via Gibson assembly, and Sanger sequenced individual colonies to determine the exact allelic sequences. Based on the obtained sequences, we designed four sgRNAs using CHOPCHOP^98^ and cloned each sgRNA into the pX330 vector (Addgene plasmid # 42230) following its standard protocol, i.e., restriction-ligation cloning of annealed sgRNA oligos into BbsI-linearized pX330 backbone. Left and right homology arms, each in 1-kb size, were also designed and PCR amplified from K562 genomic DNA. Two intermediate plasmids were constructed prior to assembling the homology directed repair (HDR) donor plasmid: first, the attP1 and attP2 gBlocks (IDT, Table S2) were inserted into a vector backbone; second, an EF1α promoter fragment and the BFP-2A-iCasp9-2A-BlastR cassette, PCR amplified from the pFL7_pLenti-pTet-Bxb1-BFP-2A-iCasp9-2A-BlastR_pCMV-rtTA3 plasmid (a kind gift from the Grimson lab),^27^ were inserted between the attP1 and attP2 sites; third, the left and right homology arms, along with the entire attP1-EF1α-BFP-2A-iCasp9-2A-BlastR-attP2 cassette, were inserted into a pUC19 vector backbone to generate the final donor plasmid. All three cloning steps were performed using Gibson assembly. The second intermediate plasmid was constructed to enable preliminary testing of Bxb1 recombination in a plasmid context prior to chromosomal integration (data not shown). Each pX330-sgRNA plasmid was then co-transfected with the donor plasmid into the ΔeNMU K562 cell line. After episomal BFP signal died out, CRISPR knock-in efficiency was assessed by gain of stable BFP expression. Three out of four sgRNAs produced a significant BFP^+^ population compared to the donor-only negative control. Single cells from these three populations were sorted into 96-well plates to derive clonal cell lines.

Outgrown single cell clones were first screened by genotyping PCR to identify those with heterozygous landing pad (LP) integration. Genomic DNA from a subset of candidate clones was purified by phenol-chloroform extraction, and a qPCR-based copy number analysis was performed by comparing BFP DNA Ct values to a control locus known to exist in three alleles in K562. Confirmed single-copy LP clones were further evaluated based on the percentage of BFP^+^ cells and Bxb1 recombination efficiency (see below). Two clonal lines, E5 and D17, were selected for subsequent experiments.

### Bxb1 recombination efficiency and eNMU rescue experiment

To construct the attB-containing payload plasmid, the attB1 and attB2 gBlocks (IDT, Table S2) were first inserted into a vector backbone to generate an intermediate plasmid. An EF1α promoter fragment and an EGFP or mCherry fragment were then introduced into the linearized intermediate plasmid. For all subsequent **individual** element cloning (i.e., excluding eNMU mutant library cloning), this parental attB1-EF1α-EGFP/mCherry-attB2 plasmid was digested with BmtI and BspEI (NEB) and the EF1α-EGFP/mCherry cassette was replaced with intended elements. All cloning steps were performed using Gibson assembly.

To evaluate Bxb1 recombination efficiency and test the functionality of the eNMU landing pad, we co-transfected the pFL9_pCAG-NLS-HA-Bxb1 plasmid (Addgene # 51271, a kind gift from the Grimson lab, transiently expressing Bxb1 recombinase) and the attB1-eNMU-attB2 payload plasmid into the LP cell lines. About 7 days post-transfection, percentage of BFP^−^ population became stable and was measured on the Sony MA900 cell sorter compared to a no-payload negative control. Both E5 and D17 LP clones consistently exhibited 4∼10% BFP loss across independent experiments, with Clone E5 showing slightly higher recombination efficiency. The recombinant BFP^−^ cells were further sorted as bulk populations and propagated for another 10∼14 days to allow stable *NMU* reactivation. Cells were then harvested for RNA extraction and RT-qPCR analysis (see below) to confirm the rescue of *NMU* gene expression.

### eNMU mutant library design, cloning, and integration

Given the central role of TFs in enhancer function, we first sought to dissect how **specific** TF binding events contribute to eNMU activity by maximizing both the extent and specificity of TF binding disruption. To this end, we aimed to (1) curate a list of motifs for TFs that are expressed in K562 cells and exhibit motif-specific binding supported by public ChIP-seq data, and (2) introduce point mutations across all motif occurrences of each selected TF. Specifically, we retrieved all available K562 ChIP-seq peaks that overlap the eNMU region (hg38 coordinate = chr4:55729891–55730846) using the UCSC Table Browser tool.^99^ We removed entries corresponding to non-sequence-specific cofactors and TFs not expressed in K562, based on ENCODE^3^ polyA plus RNA-seq data (accession: ENCSR000CPH) using a TPM > 1 threshold. Binding motifs for the remaining TFs were then obtained from the JASPAR 2022 database^29^ with few occasions from the cis-BP database^100^ (see Table S1). These motifs were further manually reviewed and filtered to retain only those located under a ChIP-seq peak. For TFs with motifs that perfectly overlap at least once—such as AP-1/NFE2 and KLF1/SP1—we grouped and treated them as a single TF. To maximize disruption of TF binding while minimizing unintended effects on adjacent motifs, we identified the two most conserved bases in each motif using position frequency matrices (PFMs) from the JASPAR 2022 database^29^ and introduced transversion mutations (A↔C, T↔G). We chose this transversion scheme because it has been shown to be more effective than alternative mutagenesis schemes.^32,33^

To complement the targeted motif mutagenesis, we designed tiling deletions across the 956-bp eNMU region, each spanning ∼100-bp intervals within the sub-elements e1 (first 453 bp) and e2 (last 503 bp). These deletions were intended to encompass GRO-cap–defined TSSs and TF motif clusters, resulting in segments e1.1–e1.4 and e2.1–e2.4. Additional segments e1.5–e1.9 were included to help resolve critical sequence features within e1.3 and e1.4. Detailed information on all mutated motifs and deleted segments is listed in Table S1.

Given the functional distinction between e1 (enhancer) and e2 (facilitator), we aimed to introduce mutations in either e1, e2, or both to dissect their individual contributions and cooperative interactions. To achieve this, we employed a “mix-and-match” cloning strategy (Figure S2D). For TF motif mutagenesis, mutant versions of e1 and e2 were synthesized separately by Twist Bioscience as dsDNA fragments, with all occurrences of a given TF’s motif mutated simultaneously. Each mutated e1 element was paired with either a wildtype e2 or a mutated e2 of the same TF type, and vice versa. Each pair was Gibson assembled with two half-backbone fragments: a fixed attB1-containing fragment and an attB2-containing fragment carrying a unique 8-bp random barcode generated by PCR. Tiling deletion constructs were built using the same cloning strategy, except that e1 deletion fragments were PCR amplified from pre-existing mutant plasmids created using the Q5 site-directed mutagenesis kit (NEB) in earlier experiments, rather than synthesized. Wildtype eNMU, Δe1, and Δe2 constructs were included as controls with known enhancer activities. Additionally, six exogenous sequences from a published STARR-seq library,^9^ kindly provided by the Yu lab, were also cloned as controls. These included the 584-bp CMV enhancer (CMV584), commonly used as a positive control in episomal enhancer reporter assays, and several non-regulatory open reading frames (ORFs), including EGFP and four human ORFs (ORF56714, ORF52920, ORF54588, and ORF55756). In total, 83 individual Gibson assembly reactions were performed and transformed into NEB Stable competent *E. coli* cells (prepared using the *Mix & Go! E. coli* transformation kit from Zymo Research).

For each Gibson assembly transformation, 8 colonies were picked and cultured overnight in deep-well 96-well plates. Colony PCR was performed on 1:20 water-diluted liquid cultures to screen for positive insertions using Q5 High-Fidelity 2X Master Mix (NEB) with primers ZZ041 and ZZ044 (Table S2), which amplify the insertion from regions flanking the Bxb1 recombination sites. The PCR program was: initial denaturation 98°C for 5 min; 30 cycles of 98°C for 10 s, 61°C for 30 s, 72°C for 39 s; and final extension 72°C for 5 min. Positive PCR amplicons (∼1.3 kb) were then purified using homebrew SPRI beads^101^ (0.7× bead ratio) and subjected to Sanger sequencing to verify element sequences and determine element-barcode associations. In total, we identified 328 unique barcodes corresponding to the 83 elements. These confirmed liquid cultures were pooled together for Maxiprep (Zymo Research) to extract the plasmid library.

Four million LP cells of the Clone E5 or D17 (biological replicates) were transfected with the plasmid library and the pFL9_pCAG-NLS-HA-Bxb1 plasmid to achieve a minimum coverage of 200× for each unique barcode representation in the recombinant population. On Day 7 post-transfection, BFP^−^ cells were sorted at a minimum coverage of 200× and expanded for another 14 days to allow full activation of *NMU*. The recombinant cells were then subjected to HCR-FlowFISH.

### HCR-FlowFISH and sequencing library preparation

To measure enhancer activity of individual elements within the pooled recombinant population, HCR-FlowFISH was performed according to the published protocol^28^ with minor modifications. We first obtained HCR probe sets and fluorescent hairpins from Molecular Instruments for the target gene *NMU* (B1 hairpin, Alexa Fluor 647 or AF647) and the internal control gene *ACTB* (B2 hairpin, Alexa Fluor 488 or AF488). Note that the *NMU* probes were custom-designed in the published study,^28^ while the *ACTB* probes were pre-designed and optimized by Molecular Instruments. FISH probing was performed in strict accordance with the published protocol,^28^ including all solution volumes and centrifugation parameters. Briefly, 20 million recombinant cells of each biological replicate were fixed with 4% formaldehyde in PBST (1× PBS, 0.1% Tween 20) at room temperature for 1 h and washed with PBST for 4 times. Following 10 min up to 24 h incubation with cold 70% Ethanol at 4°C, cells were washed with PBST twice and incubated with the pre-warmed Probe Hybridization Buffer at 37°C for 30 min. HCR probes for *NMU* and *ACTB* were added together to cells to reach a final concentration of 4 nM per probe. The samples were then incubated overnight with agitation in a 37°C hybridization oven. On the next day, cells were washed with the Probe Wash Buffer for 5 times, with 5× SSCT (5× SSC, 0.1% Tween 20) once, and pre-amplified in the Amplification Buffer for 30 min at room temperature. Snap-cooled hairpins were diluted in the Amplification Buffer and then added to the pre-amplified samples to reach a final hairpin concentration of 60 nM. Samples were incubated with rotation in a dark room overnight at room temperature. On the next day, 5× volume of 5× SSCT was added to the samples before centrifugation and removal of the hairpin amplification solution. Cells were then washed with 5× SSCT for 6 times before final resuspension in PBS at a density of 10 million cells/mL. The samples were filtered through a 35 µm Cell Strainer cap into a 5 mL polystyrene tube (Corning) before sorting.

Cells were sorted into 8 bins (2 rounds of 4-way sorting) based on the AF647/AF488 ratio (Figure S2E) at a minimum coverage of 500× barcode coverage per bin to ensure robust representation in the sequencing library. Sorted cells, together with the unsorted background sample, were pelleted and resuspended in 400 µL of ChIP lysis buffer (50 mM Tris-HCl, pH 8, 10 mM EDTA, 1% SDS), and de-crosslinked overnight at 65°C with 1000× rpm shaking. Samples were then treated with RNase A (Thermo Scientific) and Proteinase K (Invitrogen) before phenol-chloroform extraction of genomic DNA (gDNA). Sequencing libraries were prepared by two rounds of PCR using Q5 High-Fidelity 2X Master Mix (NEB). The 1^st^ round PCR was performed on the recovered gDNA corresponding to a minimum of 200× barcode coverage, using primers ZZ145 and ZZ146 (Table S2) to specifically amplify the 8-bp barcodes from the eNMU genomic locus. A maximum of 500 ng gDNA was used as input in a 50 µL PCR reaction. The PCR program was: initial denaturation 98 °C for 3 min; 11 cycles of 98°C for 10 s, 65°C for 30 s, 72°C for 1 min; and final extension 72°C for 5 min. The PCR products were then purified using homebrew SPRI beads (1.5× bead ratio) to remove unused primers, followed by the 2^nd^ round PCR with standard Illumina Nextera primers to append sequencing library indices and flow cell adaptors to the amplicons. The PCR program was: initial denaturation 98°C for 30 s; 11 cycles of 98°C for 10 s, 67°C for 30 s, 72°C for 20 s; and final extension 72°C for 5 min. Final PCR products were purified using the MinElute PCR purification kit (Qiagen) and DNA concentration was measured by the Qubit dsDNA High Sensitivity assay (Thermo Fisher). The libraries were pooled for sequencing on the Element Biosciences AVITI platform (2 × 80 bp paired-end sequencing).

### Individual element testing at the eNMU landing pad

For downstream functional analysis, critical eNMU mutants identified in the FlowFISH screen were cloned into the attB1-attB2 plasmid backbone without any element barcode. Several additional related mutants were designed and generated, whose sequences are listed in Table S1. These elements were integrated individually into the eNMU landing pad, and the BFP^−^ recombinants were sorted as single cells into 96-well plates to establish clonal cell lines as independent biological replicates. Three to four weeks after sorting, cells expanded to sufficient numbers for crude gDNA extraction^15^ and genotyping PCR to confirm element insertion: briefly, ∼0.2 million cells were pelleted and concentrated in 20 µL of culture media in a 0.5-mL PCR tube, mixed with 40 µL of Quick Extract buffer (10 mM Tris-HCl, pH 8.5, 0.45% Tween-20, 4 mg/mL protease K), and incubated at 65°C for 6 min and 98 °C for 2 min; 1 µL of the crude gDNA extract was used as input in the genotyping PCR following the standard Phusion polymerase protocol (NEB) with homemade Phusion polymerase and primers ZZ104 and ZZ105, which amplify the insertion from regions flanking the Bxb1 recombination sites. The PCR program was: initial denaturation 98°C for 3 min; 30 cycles of 98°C for 10 s, 58°C for 30 s, 72°C for 30 s; and final extension 72°C for 5 min. Positive amplicons (1132 bp) were purified using homebrew SPRI beads (0.7× bead ratio) and verified by Sanger sequencing to confirm sequence integrity. Of note, no mutations were observed in any of the single cell-derived clones, demonstrating the genomic stability of K562 cells. The verified clonal lines were subjected to RT-qPCR analysis to measure their *NMU* expression levels.

For heterologous enhancer testing at the eNMU locus, we selected candidate elements from a previously curated list of CRISPR-validated distal regulatory elements in K562 cells (Supplementary Table 6a of Fulco and Nasser et al.),^16^ prioritizing those with large effect sizes. The selected elements were PCR amplified from K562 gDNA and inserted with or without the e2 element into the attB1-attB2 plasmid backbone. Recombinant cells were sorted as bulk populations to measure enhancer activity by RT-qPCR. As a side note, the parental LP cell lines exhibited a basal BFP^−^ fraction (0.6∼2%), meaning the sorted BFP⁻ population included some non-recombinant LP cells. To estimate the true recombinant fraction, we subtracted the %BFP⁻ in the no-payload control from that in the Bxb1+payload transfection and divided this number by the total %BFP⁻ in the Bxb1+payload transfection. *NMU* expression measured by RT-qPCR was corrected based on this estimated true recombinant fraction for each element integration.

### Quantitative reverse transcription polymerase chain reaction (RT-qPCR)

K562 cells were lysed with the TRIzol Reagent (Invitrogen), and RNA was isolated using the Direct-zol RNA miniprep kit (Zymo Research) with 15 min DNase treatment on column. Reverse transcription was performed using M-MuLV RT (NEB M0253L) and Random Primer Mix (NEB S1330S) following manufacturer’s instructions. Real-time quantitative PCR (qPCR) was carried out with a custom protocol: 1/10 volume of cDNA, 1× Phusion HF Buffer (NEB), 500 nM of each primer, 200 μM dNTPs (Thermo Fisher), 0.7× SYBR Green I (Invitrogen), and 1/100 dilution of homemade Phusion polymerase. All qPCR reactions were run in technical triplicates in 10 μL volumes in 384-well plates on a Roche LightCycler 480 Instrument II with the following program setting: initial denaturation 98°C for 2 min; 45 cycles of 98°C for 10 s, 58°C for 20 s and 72°C for 30 s; melt curve 98°C for 5 s, 55°C for 1 min, ramp to 98°C at 0.11°C/s; and cool down to 40°C. *NMU* expression was normalized to the housekeeping gene *ACTB* using the 2-ΔΔCT method.^102^ Primers used for RT-qPCR are listed in Table S2.

### Chromatin Immunoprecipitation (ChIP)

ChIP experiments were conducted using two independently derived single cell clones as biological replicates for each of the four genotypes of interest: WT_eNMU, e1_mGATA1, e1_mRUNX1, e2_mSTAT5. For GATA1 and RUNX1 ChIP, cells were washed twice with ice-cold PBS and crosslinked with 1% formaldehyde (Electron Microscopy Sciences) at room temperature for 10 min before quenching by 200 mM glycine at room temperature for 5 min. For STAT5 and p300 ChIP, cells were first crosslinked with 2 mM disuccinimidyl glutarate (Santa Cruz) at room temperature for 30 min, washed 3 times with PBS and then crosslinked with 1% formaldehyde at room temperature for 5 min before quenching with 200 mM glycine at room temperature for 5 min. Two additional PBS washes were performed, and cell pellets were lysed with Farnham Lysis Buffer (5 mM PIPES, pH 8, 85 mM KCl, 0.5% NP40, 10 mM glycine, 1× Thermo Scientific Pierce Protease Inhibitor) on ice for 20 min. After centrifugation and supernatant removal, the nuclear pellet was resuspended in RIPA Lysis Buffer (10 mM Tris-HCl, pH 8, 150 mM NaCl, 1 mM EDTA, 1% NP-40, 0.5% sodium deoxycholate, 0.1% SDS, 1× Thermo Scientific Pierce Protease Inhibitor) and incubated on ice for 10 min. Sonication was carried out using a Diagenode Bioruptor device at High Setting, 30 sec on/30 sec off for three rounds of 10-min cycle to shear chromatin to a size of 100∼300 bp. The lysate was then clarified by centrifugation at 20,000 r.c.f., 4°C for 15 min, of which 2% was kept as ChIP input.

The following antibodies were used for IP: GATA1, Abcam ab11852; RUNX1, Abcam ab23980; STAT5, R&D Systems AF2168; p300, Abcam ab14984; normal rabbit IgG control, Cell Signaling Technology 2729S; normal mouse IgG1 control, Santa Cruz sc-3877. Each IP used 4 million cells/4 µg antibody/40 µL Dynabeads Protein A (for rabbit IgG) or Protein G (for mouse IgG1) (Thermo Scientific). Beads were washed three times with 5 mg/mL BSA in PBS and incubated with corresponding antibodies at 4°C for 6 h to overnight with rotation. Another three BSA/PBS washes were performed to remove unbound antibodies, and the clarified chromatin lysate was added to the beads and incubated overnight at 4°C with rotation. Beads were then washed with the following buffers, each for three times: Low Salt Wash Buffer (20 mM Tri-HCl, pH 8, 2 mM EDTA, 150 mM NaCl, 1% Triton X-100, 0.1% SDS), High Salt Wash Buffer (20 mM Tri-HCl, pH 8, 2 mM EDTA, 500 mM NaCl, 1% Triton X-100, 0.1% SDS), and LiCl Wash Buffer (10 mM Tri-HCl, pH 8, 1 mM EDTA, 250 mM LiCl, 1% NP-40, 1% sodium deoxycholate). After one final wash with TE Buffer (10 mM Tris-Cl, pH 8, 1 mM EDTA), chromatin was eluted from beads by two rounds of incubation with ChIP elution buffer (1% SDS, 0.1 M sodium bicarbonate). Each incubation involved 15 min shaking at 1200 rpm at 65°C, followed by 15 min rotation at room temperature. The eluates and input samples were treated with RNase A at 37°C for 30 min, de-crosslinked at 65°C overnight with 900 rpm shaking, followed by Proteinase K treatment at 45°C for 2 h with shaking. DNA was purified using MinElute PCR Purification Kit (Qiagen).

For GATA1, RUNX1 and STAT5 ChIP, qPCR was performed on the purified input and eluate samples to measure enrichment of TF binding at the eNMU locus. A negative control locus was also probed to estimate background signal of non-specific pull-down. Primers used for ChIP-qPCR are listed in Table S2. Since ChIP eluates were in low abundance and could be difficult to quantitate, qPCR was carried out with a custom 10× reaction mix from previous studies^103,104^ that gave great sensitivity and specificity. The 10× reaction mix composition was: 400 mM 2-amino-2-methyl-1,3-propanediol (pH adjusted to 8.3 using HCl), 50 mM KCl, 30 mM MgCl_2_, 0.09% Brij C10, 0.15% Brij 58, 500 μg/mL BSA, 300 μM dNTPs, 16.24% glycerol (v/v), 1/3000 SYBR Green I (10,000× stock), and 0.4 U/μL Platinum *Taq* DNA polymerase (Invitrogen). Final ChIP-qPCR condition was optimized to be: 1/10 volume of ChIP material, 1× custom reaction mix, 500 nM of each primer, 170 μM dNTPs, 0.35× SYBR Green I. All qPCR reactions were run in technical triplicates in 10 μL volumes in 384-well plates on a Roche LightCycler 480 Instrument II with the following program setting: initial denaturation 95°C for 10 min; 45 cycles of 95°C for 10 s, 60°C for 8 s and 72°C for 14 s; melt curve 95°C for 5 s, 45°C for 30 s, ramp to 95°C at 0.11°C/s; and cool down to 40°C. Serial dilutions of one input sample was included in each qPCR run to generate standard curves for each primer set, from which the amplification efficiency (E) was calculated. ChIP enrichment was then determined as percent input using the following equation:

%Input = 100% × E^(Ct_input − Ct_ChIP) × Input Fraction

Where E represents the qPCR amplification efficiency (ranging from 1.8 to 2.0 in our experiments), and Input Fraction refers to the proportion of total chromatin lysate used for the input (2%, or 0.02, in our case).

For p300 ChIP, sequencing libraries were prepared using a Tn5 tagmentation-based protocol. Briefly, 20 μL tagmentation reactions were set up with 0.35 ng of ChIP DNA or 1 ng of input DNA, 1× TAPS-DMF Buffer (10 mM TAPS-NaOH, pH 8.5, 5 mM MgCl_2_, 10% DMF), and 1 μL of 1:15 diluted homemade Tn5 transposase^105^ (a kind gift from Dr. Roman Spektor). The reaction was incubated at 55°C for 10 min, and 2 μL of 1% SDS was added immediately, followed by another 55°C incubation for 7 min to strip off Tn5 from DNA. Post-tagmentation PCR was carried out in 100 μL volume with 10 μL of the tagmentation reaction, 1× Phusion HF Buffer, 200 μM dNTPs, 400 nM of each Illumina Nextera index primer, and 1 μL of homemade Phusion polymerase. The PCR program was: initial extension 72°C for 3 min; initial denaturation 98°C 30s; 13 cycles of 98°C for 10 s, 63°C for 30 s, 72°C for 3 min; and final extension 72°C for 5 min. PCR products were purified first using the MinElute PCR purification kit (Qiagen), followed by an additional cleanup with homemade SPRI beads (1.5× bead ratio) to ensure complete removal of unused primers. DNA concentration was measured by the Qubit dsDNA High Sensitivity assay (Thermo Fisher). The libraries were pooled for sequencing on the Element Biosciences AVITI platform (2 × 80 bp paired-end sequencing).

### ATAC-seq

For TF motif mutants, ATAC-seq was performed on two independently derived recombinant single cell clones as biological replicates. For WT K562 and CRISPR deletion cell lines (ΔeNMU, Δe1, and Δe2) obtained from the Yu lab,^9^ ATAC-seq was conducted on two independent cultures from the same clonal source, as only one deletion clone was available for the genotypes ΔeNMU and Δe1. ATAC-seq was performed on 50,000 K562 cells with all buffer compositions and reaction conditions following the published Omni-ATAC protocol^106^ unless otherwise specified. Briefly, cells were pelleted, washed once with ice-cold PBS, resuspended in ice-cold Lysis Buffer, and incubated on ice for 3 min. Upon addition of Wash Buffer and gentle inversion, nuclei were pelleted, and supernatant (cytoplasm) was discarded. Nuclei were then resuspended gently in 50 μL transposition reaction mix containing 1 μL homemade Tn5 transposase and incubated at 37°C for 30 min with 1000 rpm shaking. DNA was purified using the MinElute PCR purification kit (Qiagen) and eluted in 21 μL volume. The entire product (∼20 μL) was mixed with 2.5 μL of each Nextera index primer (25 μM) and 25 μL NEBNext High-Fidelity 2X PCR Master Mix and subjected to a first round of PCR: initial extension 72°C for 5 min; initial denaturation 98°C 30s; 5 cycles of 98°C for 10 s, 63°C for 30 s, 72°C for 1 min. To determine additional cycles needed to avoid over-amplification, a qPCR analysis was performed using 5 μL of the first-round PCR reaction.^107^ For all ATAC-seq libraries, 3∼4 additional cycles of amplification were performed on the remaining 45 μL PCR reaction, followed by sequential cleanup using the MinElute PCR purification kit (Qiagen) and homebrew SPRI beads (1.5× bead ratio). DNA concentration was measured by the Qubit dsDNA High Sensitivity assay (Thermo Fisher). The libraries were pooled for sequencing on the Element Biosciences AVITI platform (2 × 80 bp paired-end sequencing) or Illumina NovaSeq X Plus platform (2 × 150 bp paired-end sequencing).

### PRO-seq

PRO-seq was performed on the same clonal lines used for ATAC-seq, as well as on two independently derived recombinant single cell clones harboring e1_WT or e1_mGATA1 integration. All buffer compositions and reaction conditions followed the published protocol of Mahat and Kwak et al.^108^ unless otherwise specified. Briefly, 5 million K562 cells were mixed with 250,000 Drosophila S2 cells (5% spike-in), pelleted at 1000 r.c.f. for 5 min at 4°C, washed once with ice-cold PBS, resuspended with ice-cold permeabilization buffer at a density of 1 million cells/mL, and incubated on ice for 5 min. Cells were washed twice with the same volume of ice-cold permeabilization buffer, and resuspended in 100 μL storage buffer before immediate nuclear run-on or flash-freezing in liquid nitrogen for long-term storage at –80°C. Prior to the run-on reaction, 40 μL Dynabeads MyOne Streptavidin C1 beads (Thermo Fisher) per sample were pre-washed sequentially with Hydrolysis Buffer (0.1N NaOH + 50 mM NaCl), High Salt Wash Buffer, and Binding Buffer. Pre-washed beads were resuspended in 60 μL Binding Buffer per sample. The nuclear run-on reaction was performed at 37°C for 5 min with a final concentration of 20 µM each of Biotin-11-CTP, Biotin-11-UTP, ATP, and GTP. Following RNA extraction by Trizol LS (Invitrogen) and RNA fragmentation by base hydrolysis, 30 μL pre-washed C1 beads and 30 μL Binding Buffer were added to the ∼60 μL RNA sample, and bead binding was performed at room temperature for 20 min on a rotational device. Beads were then washed twice using 500 μL High Salt Wash Buffer and once using 500 μL Low Salt Wash Buffer, with tube swap after each wash. Biotinylated RNA was eluted from beads by Trizol extraction, and 3′ RNA adaptor ligation was performed in a total volume of 20 μL with 5 µM final adaptor concentration and 2 μL T4 RNA Ligase I, High Concentration (NEB). The reaction was incubated at 20°C for 4 h and held at 4°C overnight. On the next day, 50 μL Binding buffer and 30 μL pre-washed C1 beads were added to the reaction, and another round of bead binding and bead washing was performed as described above. Subsequent 5′ enzymatic modifications of RNA were performed on beads with a reaction volume of 20 μL assuming 1 μL bead volume: 5′ decapping reaction involved 1 μL RppH (NEB) and 1-h incubation at 37°C; 5′ hydroxyl repair involved 1 μL T4 PNK (NEB) and 1-h incubation at 37°C. Beads were then washed once with 300 μL Binding Buffer, and 5′ RNA adaptor ligation was performed on beads in a total volume of 20 μL with 5 µM final adaptor concentration and 2 μL T4 RNA Ligase I, High Concentration (NEB), incubated at room temperature for 1 h on a rotational device. Beads were washed twice using 500 μL High Salt Wash Buffer and once using 500 μL Low Salt Wash Buffer, with tube swap after each wash. RNA was eluted from beads by Trizol extraction and resuspended in 13 μL RT resuspension mix (8 μL DEPC H_2_O, 4 μL of 10 μM Illumina RP1 primer, 1 μL of 10 mM dNTPs). RNA was denatured at 65°C for 5 min and snap cooled on ice, and 7 μL RT master mix was added to the sample (4 μL of 5× RT Buffer, 1 μL of 100 mM DTT, 1 μL Invitrogen SUPERase·In RNase Inhibitor, 1 μL Thermo Scientific Maxima H Minus Reverse Transcriptase). RT reaction program was: 50°C for 30 min, 65°C for 15 min, 85°C for 5 min, hold at 4°C. The resulting cDNA was diluted with an equal volume of DEPC H_2_O. A test amplification was performed on 1:4 serial dilutions of 2 μL cDNA sample and run on 6% native PAGE TBE gel to determine the optimal PCR cycle number (N). Final amplification was performed in 100 μL volume (32.5 μL DEPC H_2_O, 20 μL of 5× Phusion HF Buffer, 20 μL of 5 M betaine, 2.5 μL of 10 µM Illumina RP1 primer, 2.5 μL of 10 µM Illumina indexing RPI-n primer, 2.5 μL of 10 mM dNTPs, 1 μL homemade Phusion polymerase, 19 μL cDNA sample). PCR program was: initial denaturation 95°C for 2 min; 5 cycles of 95°C for 30 s, 56°C for 30 s, 72°C for 30 s; N cycles of 95°C for 30 s, 65°C for 30 s, 72°C for 30 s; final extension 72°C for 5 min. PCR products were sequentially purified using the MinElute PCR purification kit (Qiagen) and homebrew SPRI beads (1.5× bead ratio) to remove all unused primers. DNA concentration was measured by the Qubit dsDNA High Sensitivity assay (Thermo Fisher). The libraries were pooled for sequencing on the Element Biosciences AVITI (2 × 80 bp paired-end sequencing) or Illumina NovaSeq 6000/X Plus platform (2 × 150 bp paired-end sequencing). Note that the 3′ and 5′ RNA adaptors contain a 6-nt unique molecular identifier (UMI) to enable accurate identification of PCR duplicates in downstream bioinformatic analysis. The complete sequences of the adaptors can be found in Judd et al.^109^

### Sequencing data analysis

**Next Generation Sequencing (NGS) data preprocessing:** For all NGS sequencing data, the quality of FASTQ files was first accessed using FastQC,^110^ and the Illumina sequencing adaptors were trimmed using fastp.^111^ Note that one of the ATAC-seq samples was sequenced at 2 × 150 bp instead of 2 × 80 bp. To ensure consistency across samples, the raw FASTQ files for this sample were trimmed to 80 bp prior to any analysis.

**Activity score calculation of HCR-FlowFISH:** First, 8-bp element barcodes were extracted from the trimmed FASTQ Read 1 using fastx_trimmer^112^ with flags -Q33 -f 23 -l 30. The FASTQ format was converted into FASTA format using seqtk^113^ with the command “seqtk seq -a -q20 -n N”. Occurrences of each barcode were counted using fastx_collapser.^112^ The 328 unique barcodes were associated with their corresponding 83 elements using a custom python script. We excluded 13 barcodes that had <50 raw reads in the unsorted background sample in at least one biological replicate. Fraction of each barcode in each bin was calculated. In parallel, the mean *NMU* fluorescence intensity (AF647) of each bin was normalized by the mean *ATCB* fluorescence intensity (AF488) of the same bin. The activity score of each barcode was then calculated using the weighted average method shown in Figure S2G. Processed barcode read counts across bins and calculated barcode activity scores are provided in Table S3.

**Multiplicative model of double mutants:** To assess whether e1+e2 double mutations (either deletions or disruptions of the same TF motif) followed a multiplicative model based on single e1 and e2 mutations (inspired by Lin et al.),^21^ we first converted the median activity scores of both single and double mutants into pseudo expression values. This conversion used the linear regression equation derived from Figure S2I, which correlates FlowFISH scores with RT-qPCR measurements of *NMU* expression in a select set of mutants. The conversion was necessary because background fluorescence in the FlowFISH assay caused the activity scores to deviate from a direct representation of gene expression levels (i.e., the regression line did not pass through the origin). Log_2_ fold changes of each mutant relative to the WT_eNMU control was then calculated using the converted pseudo expression values. For each e1+e2 double mutant, the log_2_ fold change was plotted against the sum of the log_2_ fold changes of the corresponding single mutants, as shown in Figure S3D.

**Genomics sequencing read alignment:** To enable accurate alignment of sequencing reads to the eNMU locus for ATAC-/ChIP-/PRO-seq data, we built a custom genome for each recombinant TF motif mutant using the reform command line tool.^114^ Each custom genome incorporated the exact mutant sequence flanked by Bxb1 recombination sites. Alignment was performed using the bowtie2 aligner^115^: for ATAC-seq and ChIP-seq, “-end-to-end --very-sensitive” mode was used, with ATAC-seq involving a subsequent step to remove mitochondrial reads using samtools view;^116^ Picard MarkDuplicates^117^ was then used for deduplication; for PRO-seq, rRNA read removal, alignment, and deduplication (with UMI-tools)^118^ were performed using the published bash script at https://github.com/JAJ256/PROseq_alignment.sh.^119^

**ATAC-seq and ChIP-seq data analysis:** To better normalize genomics datasets across different clonal lines, we applied a “reads-under-peaks” approach to calculate scaling factors for each sample. ATAC-seq peaks were called for each sample using HMMRATAC^120^ in the MACS3 software^121^ with “-u 90 -l 30 -c 10” parameters. To create a unified peak set for count normalization, we first identified reciprocal >50% overlaps between peaks from biological replicates using bedtools intersect “-f 0.50 -r”.^122^ Consensus peak sets from each group of replicates were then merged to create a union peak set. ChIP-seq peaks were called for each biological replicate and for pooled replicates using MACS2^121^ with the input sample as control and the parameters “--broad --broad-cutoff 0.05 --keep-dup all”. Consensus peak sets from each group of replicates was generated using a published bash script (Additional file 5 of Reske et al.),^123^ which identifies pooled peaks that show >50% reciprocal overlap with each biological replicate. These consensus peak sets were then merged to create a union peak set.

To generate the read count matrix for ATAC-seq and ChIP-seq datasets, read pairs overlapping union peaks were quantified using featureCounts.^124^ DESeq2^125^ was then used to calculate scaling factors for normalization. bamCoverage^126^ was used to generate normalized bigwig files at a bin size of 1 bp for ATAC-seq and 50 bp for ChIP-seq. p300 ChIP-seq signal at the eNMU locus was quantified using bigWigAverageOverBed from the kentUtils of the UCSC Genome Browser.^127^ Data plotted in the main figures are merged bigwig files from two biological replicates generated also using kentUtils of the UCSC Genome Browser.^127^

**PRO-seq data analysis:** Unnormalized 3′-end bigwigs files for PRO-seq were first generated from bam alignment files by PINTS^10^ using the option “pints_visualizer -e R1_5 --reverse-complement”. To calculate scaling factors for PRO-seq data, a collapsed list of all GENCODE v46 transcripts^128^ was first generated using the reduceByGene() function in the BRGenomics R package.^129^ The unnormalized bigwig files were loaded into Rstudio, and a DESeq2 object was generated for the combined list of all PRO-seq samples using the getDESeqDataSet() function in BRGenomics,^129^ using the collapsed transcript list to specify genomic regions of interest. Scaling factors calculated by DESeq2^125^ were applied to each bigwig file using the applyNFsGRanges() function in BRGenomics^129^ before data export. Unnormalized 5′-end bigwigs files for PRO-seq were generated from bam alignment files by PINTS^10^ using the option “pints_visualizer -e R2_5” and normalized using the same scaling factors calculated from the 3′-end bigwigs files. Normalized bigwig files of two biological replicates were merged using kentUtils of the UCSC Genome Browser for visualization.^127^

Pausing index of the *NMU* gene was calculated using the getPausingIndices() function in BRGenomics,^129^ with the promoter region defined as TSS to TSS+250 bp and the gene body region as TSS+500 bp to TES–500 bp (TSS, transcription start site; TES, transcription end site) using the promoters() and genebodies() functions in BRGenomics.^129^ Pausing indices for both individual samples and merged biological replicates were calculated and plotted in the main figures.

**Public sequencing data analysis:** All publicly available datasets visualized in this study are listed in Table S4. All ChIP-seq datasets, except for the one targeting LDB1^44^ (GEO accession: GSE142227), were obtained from the ENCODE data portal^130^ (https://www.encodeproject.org/). To ensure consistency, we re-analyzed the LDB1 ChIP-seq raw FASTQ files using the standard ENCODE ChIP-seq pipeline^131^ (https://github.com/ENCODE-DCC/chip-seq-pipeline2). For RNA-seq datasets from Li et al.,^64^ raw read counts were downloaded from GEO accession GSE214809. For RNA-seq datasets from Schulz et al.^66^ and An et al.,^65^ NCBI-generated RNA-seq raw read counts^132^ were downloaded from GEO accessions GSE128268 and GSE53983. Differential gene expression analysis was performed using DESeq2^125^ in Rstudio.

**Data visualization:** Genome browser tracks were visualized using pyGenomeTracks.^133^ All bar plots, box plots, scatter plots, line plots, volcano plots, and correlation analyses were generated using the ggplot2 package^134^ in R (version 4.2.3) and RStudio. Flow cytometry data was analyzed and plotted using FlowJo (version 10.10.0). Schematic illustrations were generated using BioRender.com under an academic license. Figures were assembled, annotated, and finalized using Adobe Illustrator. All plots used consistent color scales for cross-comparison.

## QUANTIFICATION AND STATISTICAL ANALYSIS

One-way ANOVA with Dunnett’s post hoc test using WT_eNMU as the control was applied to ChIP-qPCR analysis. The statistical details of each experiment, including exact sample sizes (n) and additional tests (e.g., Pearson’s correlation coefficient r for linear relationships), are provided in the figure legends and/or shown graphically in the figures. For RNA-seq analysis, DESeq2 was used to determine the significance of differentially expressed genes.

## SUPPLEMENTAL INFORMATION

Table S1. Sequences of all mutants analyzed in this study. Related to Figures 2, S3, 3, and 7.

Table S2. Sequences of all oligonucleotides and dsDNA fragments synthesized for this study. Related to Methods.

Table S3. Raw read counts and calculated activity scores for individual barcodes from the HCR-FlowFISH screen. Related to Figures 2, S2, and S3.

Table S4. Public datasets and their sources used in this study. Related to Figures 1, 3, S5, 6, S6, and S7.

